# The bZIP transcription factor PnAda1 functions as a regulator of virulence, fungicide tolerance and necrotrophy in the wheat pathogen *Parastagonospora nodorum*

**DOI:** 10.64898/2026.08.18.745654

**Authors:** Shota Morikawa, Leon Lenzo, Keshara Colomba Thanthrige, Steven Chang, Kar-Chun Tan, Callum Verdonk

## Abstract

Ada1 (All Development Altered-1) is a conserved but poorly characterised basic leucine zipper (bZIP) transcription factor found throughout filamentous fungi. In the wheat pathogen *Parastagonospora nodorum*, PnAda1 is required for full virulence and is transcriptionally associated with the virulence regulator PnPf2, but its biological functions remain unclear. Here, we combined comparative RNA sequencing with targeted phenotypic analyses to define the role of PnAda1 during vegetative growth and host infection. Deletion of *PnAda1* did not abolish pathogenicity but delayed disease progression, with the *PnAda1*-deletion mutant transcriptome at 7 days post-inoculation resembling that of the wildtype SN15 at 3 days. This developmental delay was associated with impaired activation of early infection-associated genes, including putative carbohydrate-active enzymes, proteases, transporters and other host-colonisation factors. In contrast, expression of major necrotrophic effector genes was not reduced and instead remained elevated during later stages of infection, indicating that PnAda1 is required for the timely progression of infection-associated transcriptional regulation rather than direct activation of effector genes. Beyond virulence, transcriptomic and phenotypic analyses revealed roles for PnAda1 in nitrogen assimilation, carbon utilisation, abiotic stress responses and fungicide sensitivity. Notably, *PnAda1* deletion increased sensitivity to succinate dehydrogenase inhibitor fungicides and reduced expression of succinate dehydrogenase subunit genes. Collectively, our findings identify PnAda1 as a broad regulator of developmental and infection-associated transitions in *P. nodorum* and expand current understanding of the transcriptional network underlying virulence, metabolism and stress adaptation in an important fungal wheat pathogen.

**IMPORTANCE:** Fungal pathogens of crop plants pose a major threat to global food security, and understanding how virulence is regulated may reveal new opportunities for disease control. In the wheat pathogen *Parastagonospora nodorum*, disease development depends on the coordinated expression of necrotrophic effectors and other infection-associated genes. The transcription factor PnPf2 is a central regulator of these virulence programs, but the downstream pathways that execute infection remain incompletely understood. Here, we show that the understudied bZIP transcription factor PnAda1 is an important downstream component of this PnPf2-regulatory network. Our findings indicate that PnAda1 coordinates processes required for successful host colonisation, including nutrient acquisition, stress tolerance and the timely deployment of infection-associated genes. By expanding our understanding of the transcriptional regulon that underpins fungal phytopathogenicity, this study provides new insight into how fungal pathogens establish disease and coordinate complex infection programs.

## INTRODUCTION

Transcription factors (TFs) are proteins that regulate gene expression to orchestrate cellular functions across domains of life. TFs harbour various DNA-binding domains that dictate TF ability to bind specific DNA sequences (Seshasayee et al., 2011, Weirauch and Hughes, 2011). Basic leucine zipper (bZIP) is a large class of TFs found in eukaryotes and is known to modulate a diverse range of functions (Amoutzias et al., 2007), including virulence in the case of phytopathogenic fungal species (John et al., 2021). Ada1 (All Development Altered-1) is a bZIP TF characterised in several fungal species, mostly through high-throughput gene knockout characterisations, and thus its function is only broadly known (**Table 1**).

**Table 1:** Characterised Ada1 orthologs and their reported functions in published literature. Rows are ordered by publication date.

| Species | Name | Locus ID | Reported Functions | Reference |
| --- | --- | --- | --- | --- |
| *Neurospora crassa* | *Ada-1* | *NCU00499* | Vegetative growth & carbon metabolism | (Tian et al., 2011) |
| *Fusarium graminearum* | *Gzbzip001* | *FGSG_00515* | Vegetative growth & virulence | (Son et al., 2011) |
| *Magnaporthe oryzae* | *MobZIP10* | *MGG_04758* | Vegetative growth, conidiation & appressorium formation | (Kong et al., 2015) |
| *Fusarium pseudograminearum* | *FgAda1* | *FPSE_04421* | Vegetative growth, virulence, conidiation & cell cycle | (Chen et al., 2020) |
| *Alternaria alternata* | *AaAda1* | *AALT_g5461* | Vegetative growth, virulence, conidiation & oxidative stress tolerance | (Gai et al., 2022) |
| *Aspergillus flavus* | *bZIP4* | *AFLA_078500* | Vegetative growth, virulence, conidiation, oxidative stress tolerance, sclerotia formation & aflatoxin production | (Zhao et al., 2022) |
| *Parastagonospora nodorum* | *PnAda1* | *JI435_044860* | Vegetative growth, virulence, pycnidiation & oxidative stress tolerance | (John et al., 2024) |

Although the "Ada1" name is also applied to a structural subunit of the SAGA transcriptional co-activator complex in fungal pathogens such as *Verticillium dahliae* (Geng et al., 2022), the Ada1 orthologs characterised here are bZIP transcription factors distinct from the SAGA-complex protein. Notably, almost all characterised Ada1 orthologs in phytopathogenic fungi are involved in virulence (Chen et al., 2020, Gai et al., 2022, Son et al., 2011, Zhao et al., 2022, John et al., 2024). The sole exception was in the rice pathogen *Magnaporthe oryzae*, although the Ada1 ortholog MobZIP10 was required for the normal formation of appressoria, specialised cells involved in host penetration (Kong et al., 2015). Thus, elucidating the regulatory network of Ada1 may further shed light on the molecular basis of host-pathogen interactions.

In the necrotrophic wheat pathogen *Parastagonospora nodorum*, causal agent of septoria nodorum blotch (SNB), the Ada1 ortholog (PnAda1) was first identified as a direct regulatory target of the zinc finger (Zn_2_Cys_6_) TF PnPf2, an important regulator of virulence (John et al., 2024, Jones et al., 2019, Rybak et al., 2017). PnPf2 also regulates the expression of necrotrophic effector (NE) genes, which are the predominant drivers of host-specific *P. nodorum* pathogenicity on wheat (McDonald et al., 2023) and function as a regulatory umbrella of other downstream transcriptional regulators including PnAda1 (John et al., 2024). Both PnAda1 and PnPf2 regulate virulence and oxidative stress tolerance in *P. nodorum* (John et al., 2024). Owing to its association with the PnPf2 regulatory network, the broader role of PnAda1 in gene regulation has not been determined and compared with PnPf2 (Jones et al., 2019).

Here, we aimed to elucidate the role of PnAda1 in NE gene expression and further characterise the developmental regulator in *P. nodorum*. Further characterisation of PnAda1 was accomplished by comparative RNA sequencing (RNA-Seq) between *P. nodorum* wildtype and a mutant with the deletion of *PnAda1* during both *in vitro* vegetative growth and *in planta* infection, followed by phenotypic characterisation guided by the transcriptomic analysis. Our transcriptomics-guided approach utilised in this study broadened the understanding of the PnPf2 regulatory network that encompasses PnAda1 and potentially provides a foundation for further characterisation of this understudied, yet important, developmental regulator of fungi.

## RESULTS

### PnAda1 promotes virulence independent of host penetration

PnAda1 is required for full virulence and development in *P. nodorum*. Previously, we observed that *P. nodorum* carrying a *PnAda1* deletion displayed reduced disease severity on wheat (John et al., 2024). Initially, we wanted to ensure PnAda1-mediated pathogenicity was not perturbed during initial host penetration. To evaluate the effectiveness of the *PnAda1-*mutants on infection, we performed a detached leaf assay (DLA) by pre-wounding wheat leaves prior to fungal inoculation as previously described (Morikawa et al., 2026).

Over a 10-day infection time course, SN15 and the *PnAda1*-complement strain *PnAda1-Comp* caused comparable disease symptoms on wheat. In contrast, each of the two independent *PnAda1*-deletion mutants, *pnada1-2* and *pnada1-15*, showed delayed virulence relative to SN15 regardless of the pre-wounding (**Figure 1A**). Therefore, host penetration is unlikely to be a causal factor in the reduced virulence of the *PnAda1*-deletion mutants. Instead, it may indicate that PnAda1 is implicated in the production or an impairment in functional coordination of virulence factors that are needed for normal plant infection.

**Figure 1:**
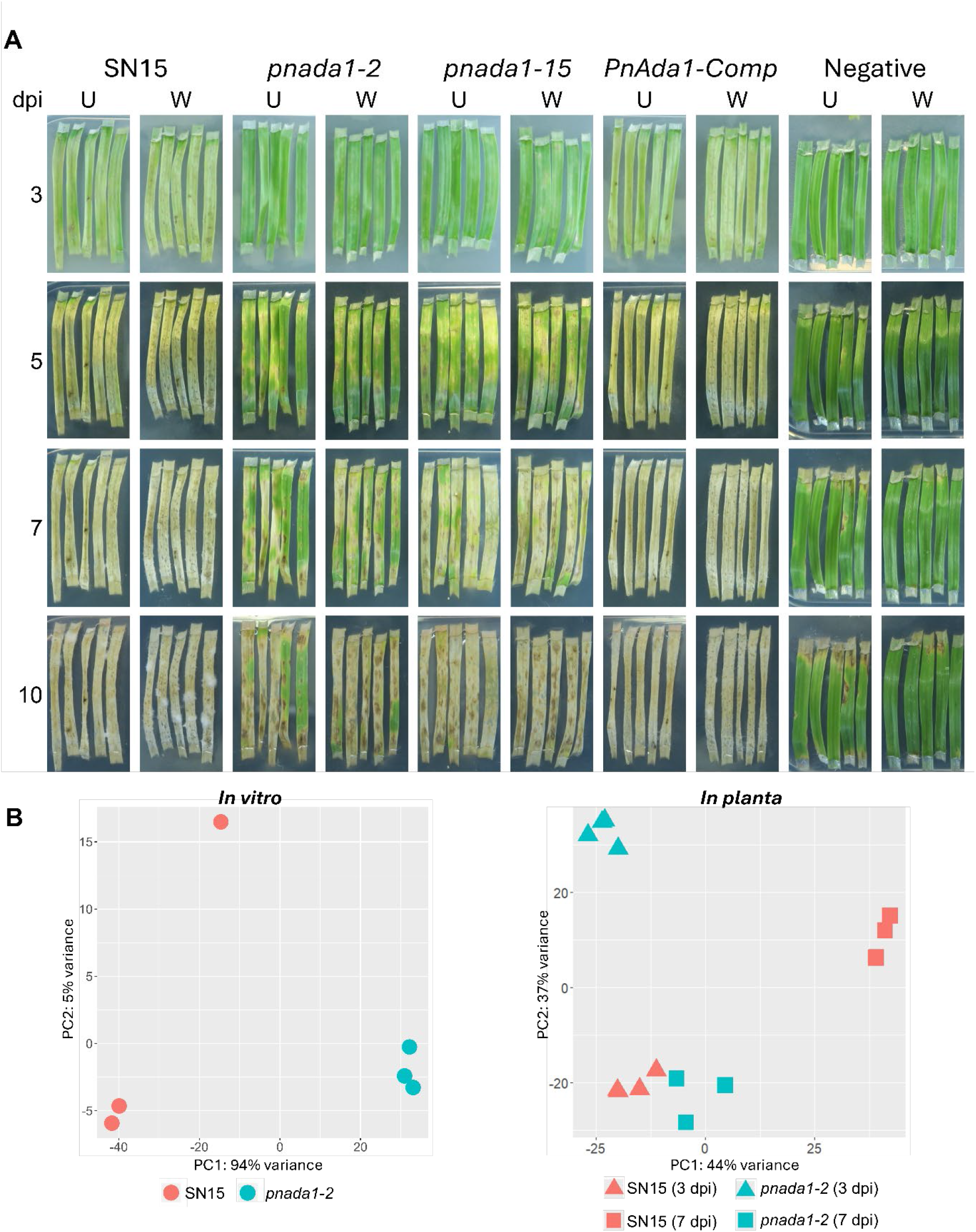
Comparison of *PnAda1*-mutants and SN15. (**A**) Detached leaf assay of *P. nodorum* SN15 wildtype, *PnAda1*-mutants *pnada1-2* and *pnada1-15,* as well as complement strain *PnAda1-Comp* infecting unwounded (U) and pre-wounded (W) wheat leaves (cv. Halberd). “dpi” – days post inoculation. (**B**) A principal components (PC) analysis of samples in the comparative RNA-Seq experiments, both *in vitro* (left) and *in planta* (right), representing a clear segregation between *P. nodorum* strains SN15 and the *PnAda1-*mutant *pnada1-2*.

### PnAda1 predominantly functions as a transcriptional repressor during axenic growth

To elucidate the potential genes associated with this observed delayed-infection phenotype, we utilised a comparative RNA-Seq approach between SN15 and the *PnAda1*-deleted mutant strain *pnada1-2* during *in-vitro* growth. A principal component (PC) analysis of the samples showed segregation between the strains (**Figure 1B**). PC1 accounted for 94% of the variation, and while one SN15 replicate segregated from the other replicates, PC2 accounted for only 5% of the variation. A scatterplot of average normalised expression of genes in SN15 against *pnada1-2* showed that high Log_2_ fold change (LFC) cutoff values deviate from the adjusted *p*-value threshold at higher normalised expression values (**Supplemental Figure S1A**). To verify whether high LFC cutoff values affect specific genes, the average normalised expression of genes in SN15 was plotted according to the genes that satisfy various LFC thresholds in *pnada1-2*. High LFC cutoffs disproportionately filter out basally highly expressed genes, and the effect was statistically more pronounced for up-regulated genes (**Supplemental Figure S1B** and **S1C**). The effect of different adjusted *p*-value thresholds was also tested, and the more stringent adjusted *p*-value thresholds expectedly filtered out genes with basal low expression, where the signal-to-noise ratio is lower (**Supplemental Figure S1D**). Therefore, an absolute LFC cutoff of > 0.58, corresponding to a 1.5-fold increase in expression for up-regulated genes, and an adjusted *p*-value threshold of <0.01 were used to define differentially expressed genes (DEGs).

Relative to SN15, we observed 2709 up-regulated and 1086 down-regulated genes in the *PnAda1-*mutant *pnada1-2* (**Supplemental Table S1**). This suggests that PnAda1 predominantly function as a transcriptional repressor *in vitro*. Out of 64 previously characterised *P. nodorum* genes for their roles on host pathogenicity, 15 were differentially expressed *in vitro*, excluding *PnAda1* itself (**Supplemental Table S2**). A Gene Ontology (GO) enrichment analysis revealed that six GO terms were over-represented in *in vitro* DEGs (**Figure 2A**). The enriched GO terms indicated up-regulation of catalysis-related and cell wall- and membrane-related genes. All enriched GO terms were up-regulated overall, but this did not translate to a statistically significant increase in medians of average gene expression between SN15 and *pnada1-2* of DEGs within enriched GO terms (**Supplemental Figure S2**). Therefore, our focus was instead given to GO terms with a high percentage of DEGs. Out of 1793 total assigned GO terms, 99 consisted of over 50% DEGs. Of this subset, 33 GO terms were associated with amino acid biosynthesis and nitrogen assimilation, further implicating *PnAda1* in nitrogen metabolism. Kyoto Encyclopedia of Genes and Genomes (KEGG) pathway enrichment analysis revealed enrichment of heterokaryon incompatibility proteins and leucine metabolism in up-regulated genes (**Figure 2B(i)**), while down-regulated genes were enriched in transporters and lysine biosynthesis.

**Figure 2:**
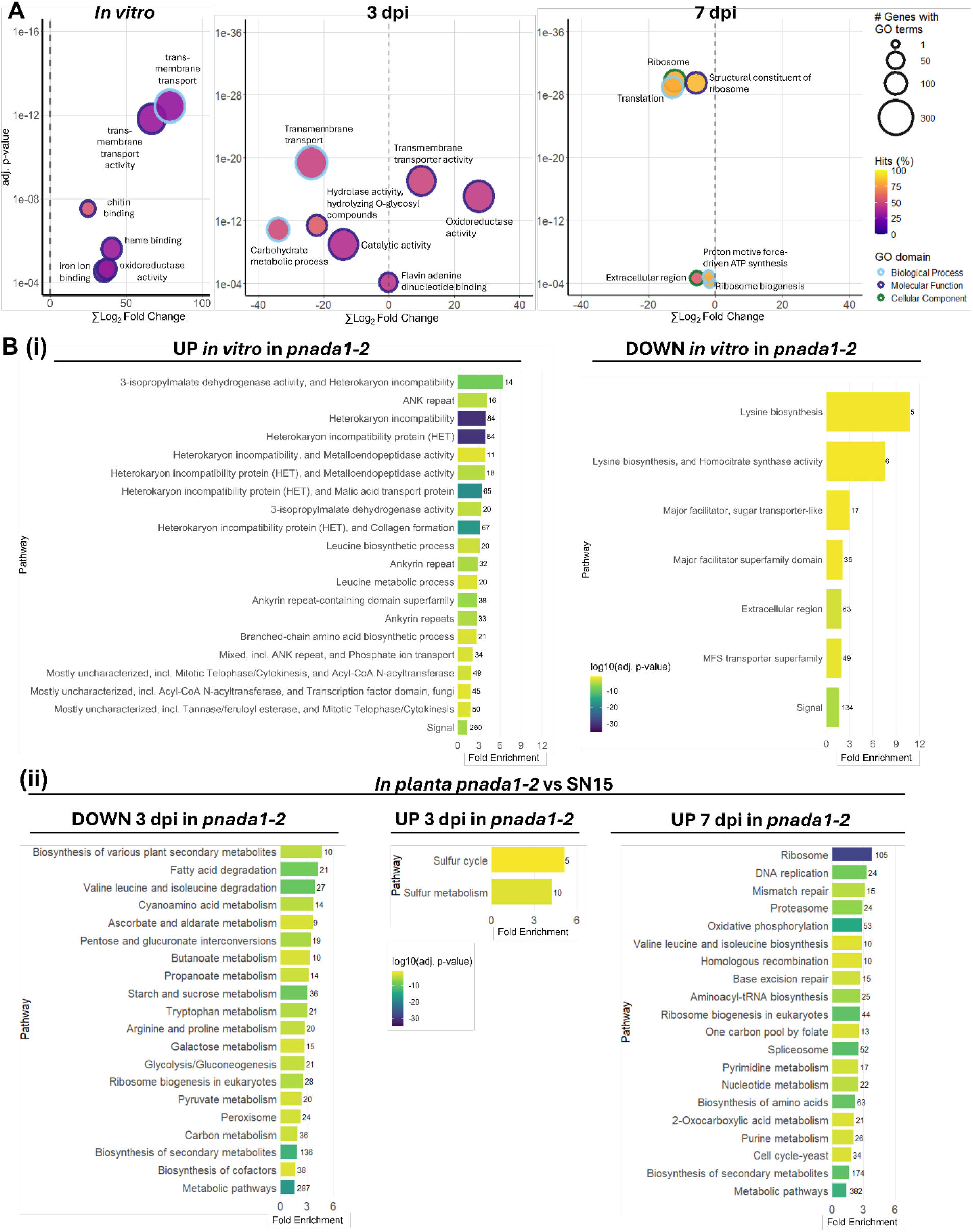
Analysis of comparative RNA-Seq between SN15 and *pnada1-2*. (**A**) Gene Ontology (GO) enrichment analysis of differentially expressed genes in *pnada1-2* relative to SN15. There is an overall up-regulation of genes involved in catalysis and encoding cell wall- and membrane-associated proteins *in vitro*, while *in planta* early infection (3 dpi) highlights the reduction of secretome-related genes and late infection (7 dpi) indicates a reduction of protein translation machinery. The x-axis indicates the sum of the LFCs of differentially expressed genes in each GO term. The y-axis indicates the adjusted *p*-value. (**B**) KEGG pathway enrichment analysis of (**i**) *in vitro* up- and down-regulated genes of *pnada1-2* relative to SN15. KEGG enrichment shows enrichment of heterokaryon incompatibility pathways in *pnada1-2* up-regulated genes, while lysine biosynthesis and transporters were enriched in *pnada1-2* down-regulated genes. (**ii**) *In planta* KEGG Up- and down-regulated genes in *pnada1-2* relative to SN15 showing a general disruption of nutrient homeostasis. There were no down-regulated genes identified in KEGG analysis for *pnada1-2* at 7 dpi. The numbers next to each bar represent the count of DEGs in each enriched pathway.

### PnAda1 deletion delays the expression of early infection genes

We then used an RNA-Seq approach to compare transcriptomes of SN15 wildtype and *pnada1-2* from infected wheat leaves *in planta* at 3- and 7-days post-inoculation (dpi). These timepoints corresponding to early host penetration and established infection, respectively (Rybak et al., 2017, Ipcho et al., 2012). PC analysis separated the samples primarily by infection timepoint and strain, with PC1 accounting for 44% of the variance and PC2 for 37% (**Figure 1B**). Notably, the *pnada1-2* transcriptome at 7 dpi clustered with SN15 3 dpi.

Here, we observed 2,410 up-regulated and 2,300 down-regulated genes in *pnada1-2* relative to SN15 at the same time-contrasts 3 dpi *in planta*. This increased to 3,524 up-regulated and 4,160 down-regulated genes at the later infection-representing 7 dpi timepoint (**Figure 3A**). The extent of differential expression *in planta* (at both 3 dpi and 7 dpi timepoints) was therefore substantially greater than that observed *in vitro* for *pnada1-2*.

**Figure 3:**
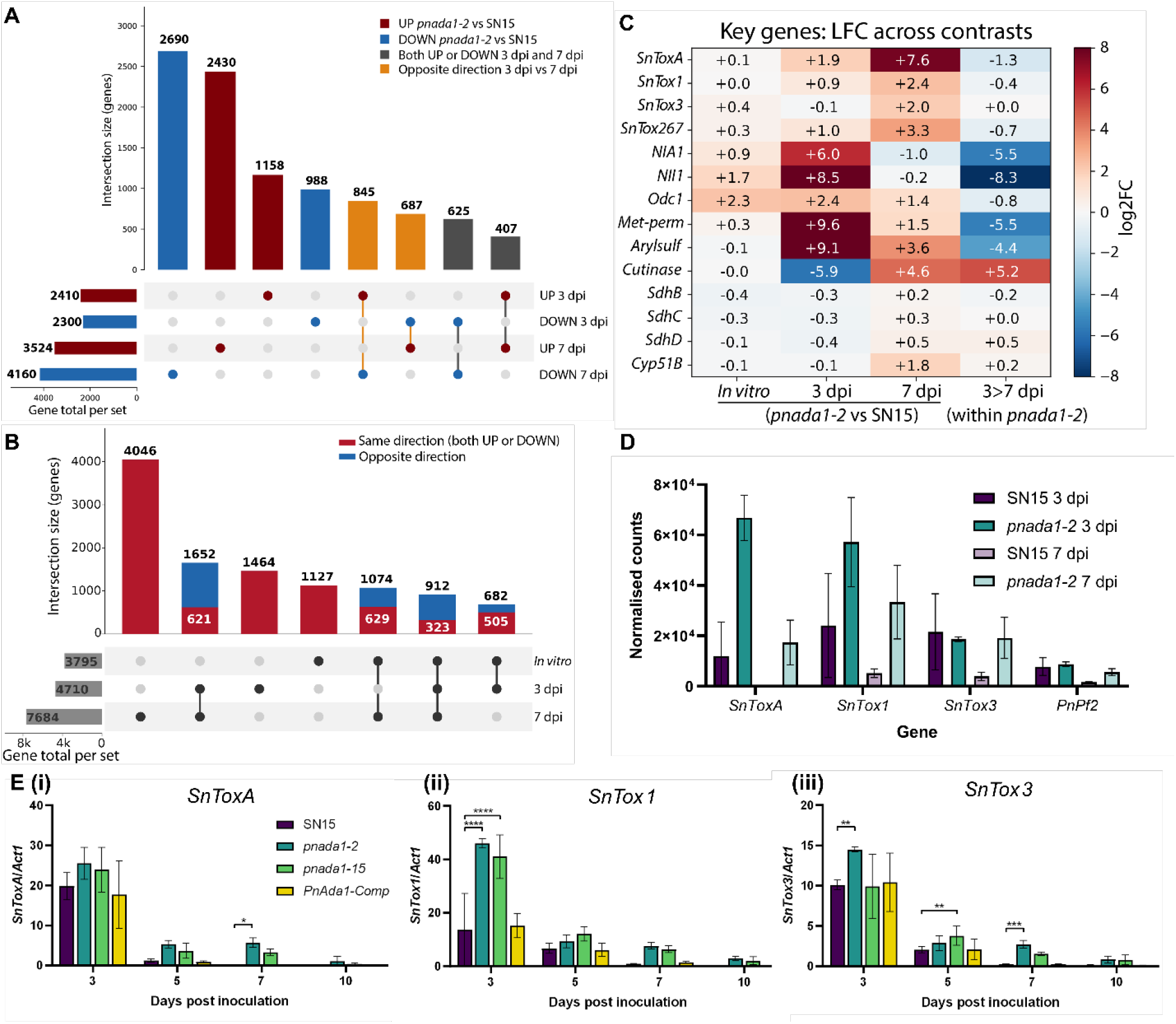
PnAda1 is implicated in key virulence gene regulation and metabolism-associated genes. (**A**) UpSet plot of DEG-set intersections across the two *in planta* timepoint contrasts, 3 dpi and 7 dpi, tested for *pnada1-2* -- highlighting total gene counts temporally regulated within *pnada1-2* relative to SN15. (**B**) UpSet plot highlighting the overlapping DEG within the *pnada1-2* RNA-Seq data across all three contrasts (*in vitro*, 3 dpi and 7 dpi *in planta*) relative to SN15 for equivalent contrasts. (**C**) Heatmap of important genes DE in the *PnAda1-*mutant determined by RNA-Seq analysis. *Met-perm* - methionine permease (JI435_146370), *Arylsulf* - arylsulfatase (JI435_020660), *Cutinase* (JI435_143050). All characterised *P. nodorum* gene differential expression can be found in **Supplemental Table S2**. (**D**) Normalised transcript read count for NE genes and the PnPf2 TF from the RNA-Seq data in SN15 and *pnada1-2*. (**E**) qRT-PCR determined gene expression profiles of necrotrophic effector genes (**i**) *SnToxA*, (**ii**) *SnTox1* and (i**ii**) *SnTox3* normalised to housekeeping gene *Act1*. Only *SnTox1* is expressed more highly in *PnAda1*-deleted mutants. Asterisks represent significant differences between SN15 and mutant strains according to a two-way ANOVA with Tukey’s HSD (*p* <0.01 (**); <0.001 (***); <0.0001 (****)).

Consistent with the PC analysis, the SN15 transcriptome remodelled greater between 3 and 7 dpi than the *pnada1-2* mutant (7,508 vs 4,193 DEGs). Indeed, 5,365 genes that changed in expression between the two timepoints (i.e. between 3 dpi and 7 dpi) in SN15 were not DE across the two timepoints in *pnada1-2*. These findings indicate that PnAda1 is required for transcriptional transition between early and late-stage establishment of infection.

To distinguish genes under consistent PnAda1-control from those reflecting delayed infection progression, DEGs were classified across all contrasts in the *pnada1-2* mutant. A core set of 323 genes were DE in the same direction across (i.e. all up or all down-regulated) all three *pnada1-2* comparisons (*in vitro*, 3 dpi and 7 dpi), comprising 205 genes repressed and 118 genes activated by PnAda1 (**Figure 3B**). This subset of genes, which account for ∼2.8% of total genes within *P. nodorum* SN15, are reflective of a common PnAda1-regulated gene set -- genes moving consistently across *in vitro* culture and both early and late-stage host-infection timepoints.

### PnAda1 is a positive regulator of CAZymes and nutrient assimilation genes

To investigate the delayed infection progression of the *PnAda1* mutants *in planta*, we interrogated the RNA-Seq data for specific causal genes with known associated functions. At *in planta* 3 dpi, genes down-regulated in *pnada1-2* were enriched for functions associated with host colonisation. Gene Ontology (GO) terms relating to the extracellular region, carbohydrate metabolic process and hydrolase activity acting on O-glycosyl compounds were over-represented among down-regulated genes (**Figure 2A**). We consistently observed down-regulation of genes that encode secreted plant cell wall-degrading enzymes, proteases and carbohydrate-active enzyme (CAZymes) in *pnada1-2*.

This includes multiple secreted serine proteases and a fungalysin-type metalloprotease (LFC = −4.7 to −5.9). Genes annotated as glycosyl hydrolases and cutinases were predominantly down-regulated in *pnada1-2* at 3 dpi (28 of 51 and 6 of 8 DEGs, respectively). Notably, this suppression was timepoint-specific: the putative cutinase *JI435_143050* was strongly down-regulated in *pnada1-2* at 3 dpi (LFC = −5.9) yet markedly up-regulated at 7 dpi (LFC = +4.6) (**Figure 3C**).

Conversely, genes up-regulated in *pnada1-2* at 3 dpi were enriched for transmembrane transporter and oxidoreductase activities. This includes pronounced induction of nutrient-scavenging genes such as a high-affinity methionine permease (*JI435_146370*, LFC = +9.6) and an arylsulfatase (*JI435_020660*, LFC = +9.1), alongside numerous major facilitator superfamily transporters. The previously characterised nitrogen assimilation genes *NIA1* and *NII1* (Cutler and Caten, 1999, Howard et al., 1999), which were highly up-regulated in *pnada1-2* at 3 dpi (LFC = +6.0 and +8.5, respectively) but not at 7 dpi (LFC = −1.0 and −0.2) (**Figure 3C, Supplemental Table S2**).

As PnPf2 also function as a positive regulator of CAZyme expression *in planta*, we then determine if there is a co-regulatory overlap between PnPf2 and PnAda1-regulated CAZymes during host infection. As PnAda1 is a direct PnPf2 target (John et al., 2024), this raises the possibility that a subset of the CAZymes attributed to PnPf2 are regulated indirectly, through PnPf2-dependent activation of PnAda1. To distinguish direct from relayed regulation, we compared the PnAda1 and PnPf2 *in planta* CAZyme regulons and scanned their promoters for the relevant *cis*-elements. Using the PnPf2-regualted CAZyme set described previously (Jones et al., 2019), we scanned all associated CAZyme genes DE in either *pnada1-2* or the PnPf2-deletion mutant *pf2ko*. Within the subset of CAZyme genes regulated by either *PnAda1* or *PnPf2*, 268 were DE within the *PnAda1*-mutant only, 11 to PnPf2 deletion only, and 126 to both PnAda1 and PnPf2 (**Table 2**). Scanning these putative CAZyme gene upstream regions’ containing promoters for the two characterised PnPf2 binding DNA motifs (M1, RWMGGVCCGA; M2, CGGCSBYWYBKCGGC) indicated all 11 of the PnPf2-only DE CAZymes were strongly enriched for the PnPf2 M1 element. The PnPf2 M1 and M2 motifs were absent (only present at genome-wide background non-significance frequency of 3.4%, less than previously observed (John et al., 2024)) in the 268 PnAda1-only regulated CAZymes. We were unable to detect any *de novo* or bZIP-like TF DNA motifs that were enriched in any of the 394 DE CAZyme genes in *pnada1-2 in planta*. Together, these observations suggest that PnPf2 and PnAda1 converge on a common early cell-wall-degradation regulon, and that PnAda1 additionally controls a larger set of CAZymes that lack PnPf2-like binding sites and are therefore unlikely to be PnPf2 targets.

**Table 2:** Shared vs distinct CAZyme regulation by PnAda1 and PnPf2 relative to SN15. PnPf2 CAZyme gene regulation described in (Jones et al., 2019). The majority of putative CAZyme genes are dictated by PnAda1 regulation, independent of PnPf2, consistent with PnPf2 activating PnAda1-expression.

| CAZyme set | Number | Fraction |
| --- | --- | --- |
| **PnAda1-regulated\*** | **394** |  |
| **PnPf2-regulated** | **137** |  |
| also PnAda1-regulated | 126 | 92% of PnPf2 CAZymes |
| PnPf2-specific | 11 | 8% of PnPf2 CAZymes |
| **Shared by both PnPf2 and PnAda1, *in planta*** |  |  |
| Same direction (both UP *or* DOWN) | 53 | 83% shared |
| Opposite direction | 11 | 17% shared |
| **PnAda1-specific *in planta*** | **268** | **68% of PnAda1 CAZymes** |

\*Across all *pnada1-2* contrasts (*in vitro*, *in planta* 3 dpi and *in planta* 7 dpi).

### *PnAda1* deletion delays host infection of wheat

We hypothesised that the deletion of PnAda1 caused an overall delay in host infection. Firstly, as described above: PCA analysis revealed that the *pnada1-2* transcriptome at 7 dpi clustered with SN15 3 dpi (**Figure 1B**). This suggests that *pnada1-2* ability to infected was severely delayed, but not arrested, compared to SN15.

Second, it was observed from the read count that the *pnada1-2* mutant produced substantially less fungal reads, suggesting lower biomass, *in planta* than SN15. This is most prominent at 3 dpi, where *pnada1-2* yielded approximately 5- to 20-fold fewer assigned fungal reads per sample (**Supplemental Figure S3**). PC1 therefore co-varied strongly with fungal read depth. The strain- and timepoint-associated separation was reproduced when the analysis was restricted to genes robustly detected across all samples, confirming a genuine transcriptional divergence between strains.

To directly validate our hypothesis that the *pnada1-2* transcriptome is developmentally delayed rather than permanently perturbed, we performed a direct comparison between DEGs for *pnada1-2* at 7 dpi against SN15 at 3 dpi. Time-matched comparison between *pnada1-2* and SN15 at 7 dpi identified 7,684 DEGs (3,524 up, 4,160 down) (**Figure 3B**), whereas the time-shifted (i.e. *pnada1-2* 7 dpi vs SN15 3 dpi) comparison identified only 3,573 DEGs (1,555 up and 2,018 down), a 54% reduction in DEGs detected. Of note, a large proponent of recovering genes within *pnada1-2* at the later 7 dpi timepoint were dominated by plant-cell-wall-active families, including GH43 (9 genes), AA9 (9 genes), GH3 (8 genes), GH31 (7 genes), AA7 (6 genes) and AA3 (5 genes). Thus, CAZymes that are associated with PnAda1 demonstrated maximal expression at 7 dpi comparable to the CAZyme transcript levels at SN15 3 dpi. This evidence further suggests a delay in deployment of virulence-factors in the *PnAda1*-deleted mutant.

### Deletion of *PnAda1* disrupted necrotrophic effector gene expression

Previous studies demonstrated that PnPf2 function as a positive regulator of *SnToxA*, *SnTox1* and *SnTox3* NE expression in *P. nodorum* (Rybak et al., 2017, Morikawa et al., 2026). As PnAda1 is a downstream target of PnPf2, we asked whether PnAda1 can also function as a co-regulator for each of these NEs. Analysis of read counts of SN15 and *pnada1-2* suggests no differences in NE gene expression *in vitro* (**Supplemental Table S2**). We then compared NE gene expression between SN15 and *pnada1-2 in planta* RNA-Seq at each of the two infection timepoints. Within SN15, expression of the NE genes *SnToxA*, *SnTox1* and *SnTox3* decreased markedly between 3 and 7 dpi, consistent with their reported early-infection expression profile (Rybak et al., 2017, Richards et al., 2022). In *pnada1-2*, this hallmark ‘high-to-low’ expression profile was not observed, thus down-regulation was abolished (**Figure 3D**). The difference was most pronounced for *SnToxA*, which was the single most up-regulated gene in the *pnada1-2* mutant at 7 dpi (LFC = +7.6; approximately 300-fold higher normalised expression than SN15), with *SnTox1* and *SnTox3* also significantly up-regulated (LFC = +2.4 and +2.0, respectively). *PnPf2* expression remained comparatively similar between SN15 and *pnada1-2* at 3 dpi, although PnPf2 transcript levels in *pnada1-2* did not fall to SN15-like levels at 7 dpi (**Figure 3D**). These findings indicate that either the contribution of PnAda1 to virulence is largely independent of NE-mediated pathogenicity, or that PnAda1 regulates other necrotrophic lifestyle genes that are needed to complement NEs for normal infection. Furthermore, PnAda1 may be required for the down-regulation of NE genes during the transition to established, late-phase infection *in planta*. To determine whether the elevated NE expression observed in *pnada1-2* at 7 dpi reflected a defect in regulation rather than sustained overexpression, we compared *pnada1-2* at 7 dpi directly with SN15 at 3 dpi. No significant differences in transcript abundance were detected for the major NEs (LFC: *SnToxA* +1.6, *SnTox1* +1.2, *SnTox3* +0.3; all *padj* > 0.01). These findings indicate that PnAda1 is required for the timely progression of NE expression during infection and supports that apparent overexpression of NE genes in *pnada1-2* reflects delayed infection development.

To validate our RNA-Seq observations for NEs, we assessed the regulation of the three key NE genes *SnToxA*, *SnTox1* and *SnTox3* in each of the *PnAda1* mutants during *in planta* host infection at all established timepoints using qRT-PCR. While variability was observed between strains, the *PnAda1*-deletion mutants expressed *SnTox1* genes comparably to our RNA-Seq data; higher than SN15 and *PnAda1-Comp* (**Figure 3E**). *SnToxA* and *SnTox3* were expressed near SN15-levels in the *PnAda1*-deletion mutants at 3 dpi, where NE gene expression is highest (Rybak et al., 2017), contrasting with our observed RNA-Seq analysis.

### PnAda1 is required for nitrogen assimilation, carbon utilisation and abiotic stress tolerance in *P. nodorum*

As KEGG pathways related to amino acid biosynthesis were enriched in DEGs, and previously characterised nitrogen metabolism genes were also differentially regulated, we investigated the involvement of PnAda1 in nitrogen metabolism. *P. nodorum* SN15, *PnAda1-Comp* and both independent *PnAda1-*mutants were grown on defined minimal media (MM) agar with varying sole nitrogen sources. *PnAda1*-deletion mutants *pnada1-2* and *pnada1-15* showed reduced growth compared to SN15 on MM supplemented with ammonium, lysine, nitrate and urea as the sole nitrogen sources (**Figure A** and **Supplemental Figure S4A**). The reduced growth of *pnada1-2* and *pnada1-15* was most drastic on MM with lysine, which is consistent with the *in vitro* transcriptomic data, as lysine metabolism genes were enriched in down-regulated genes of *pnada1-2*. Overall, PnAda1 is required for the nitrogen assimilation needed for growth.

Next, we wondered whether the reduced carbohydrate metabolic processes identified in GO enrichment *in planta* impacted *PnAda1*-dependent carbon utilisation. As above, we investigated the *PnAda1-*mutants on defined minimal media carrying different sole carbon sources and assessed their vegetative phenotype growth. *pnada1-2* and *pnada1-15* showed reduced growth compared to SN15 on media with starch (**Figure 4B** and **Supplemental Figure S4B**), consistent with the reduced CAZyme and alpha-glucosidases expression observed in the *pnada1-2* RNA-Seq. In contrast, the *PnAda1-*mutants had relatively higher growth in both sodium acetate and ethanol-containing media, suggesting that PnAda1 may limit acetate-assimilation flux.

**Figure 4:**
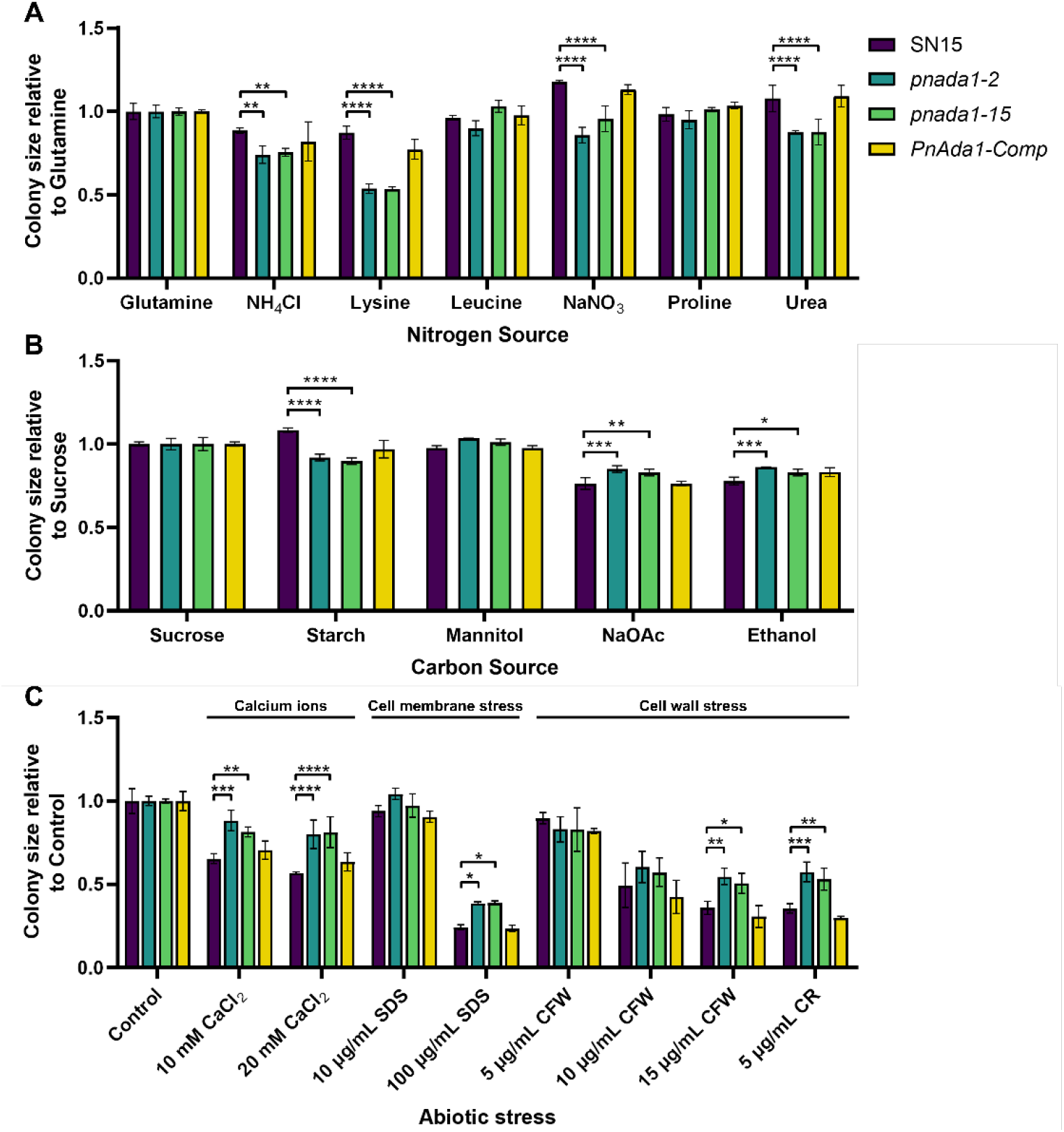
Nitrogen utilisation, Carbon metabolism and abiotic stressors differ between wildtype SN15 and *PnAda1*-deletion mutants. (**A**) Average relative (to glutamine) colony diameter for *PnAda1*-deletion mutants, SN15 and complement strain *PnAda1-Comp* grown on defined minimal media supplemented with variable sole nitrogen sources. (**B**) Average sucrose-relative colony diameter for each tested strain against carbon sources. (**C**) *PnAda1*-deletion mutants exhibit enhanced tolerance to abiotic stressors. The bar graph shows the average colony size of strains on MM agar with various abiotic stressors (identified above), normalised to a control without supplement. SDS – sodium dodecyl sulphate; CFW – Calcofluor White; CR – Congo Red. Asterisks represent significant differences between SN15 and mutant strains according to a two-way ANOVA with Tukey’s HSD (*p* <0.05 (*); <0.01 (**); <0.001 (***), <0.0001 (****)).

PnAda1 is required for H_2_O_2_ oxidative stress tolerance (John et al., 2024). DEG suggests PnAda1 possesses a role in cell wall/membrane function and repression of the calcium-signalling transcription factor gene *PnCrz1*. PnCrz1 has an involvement in oxidative stress response based on phenotypes observed during hydrogen peroxide media-containing culturing (Choupannejad et al., 2025). We then challenged the *PnAda1*-deletion mutants with excess calcium ions and cell wall/membrane stressors on defined MM to determine if these stressors antagonise vegetative growth. The *PnAda1*-deletion mutants showed increased tolerance to high calcium ion concentrations, the fungal cell wall stressors Congo Red and Calcofluor White, and sodium dodecyl sulphate, a cell membrane stressor (**Figure 4C**). These phenotypes are consistent with our transmembrane efflux-related and oxidoreductase activity GO enrichment within the *in vitro* transcriptomic data and indicate that PnAda1 represses tolerance to several abiotic stressors.

### Deletion of *PnAda1* results in hypersensitivity to SDHI fungicides

Succinate dehydrogenase subunits SdhB, SdhC and SdhD are the targets of the succinate dehydrogenase inhibitor (SDHI) class of fungicides (Duarte Hospital et al., 2023). The identified orthologs for genes encoding the subunits of the succinate dehydrogenase *SdhB* (*JI435_033510*) and *SdhC* (*JI435_111570*) exhibited a statistically significant reduction in *in vitro* expression in *pnada1-2* compared to SN15 (**Figure 5A**). There were no statistically significant changes in the expression of the *SdhA* (*JI435_165470*) or *SdhD* orthologs (*JI435_116200*), nor the previously characterised gene *Cyp51B* (Zulak et al., 2025), which encodes the essential enzyme targeted by the demethylase inhibitor (DMI) class of fungicides. However, the reduced expression of *SdhB* and *SdhC* fell below the LFC threshold as the reduction was only 20-30% of basal expression. Nevertheless, the structurally distinct SDHI fungicides fluxapyroxad and boscalid and DMI fungicides epoxiconazole and tebuconazole were used to test the SDHI and DMI sensitivity of the *PnAda1* mutants (**Figure 5B**). Increased sensitivity to fluxapyroxad and boscalid was observed in the *PnAda1-*deleted mutants at higher tested concentrations compared to SN15 (**Figure 5C**). Conversely, *pnada1-2* and *pnada1-15* exhibited sensitivity to epoxiconazole and tebuconazole comparable to SN15 (**Figure 5D**). We concluded that *PnAda1*-deletion mutants had higher-selective sensitivity to SDHI fungicides, rather than an overall increased sensitivity to all broad-spectrum fungicides.

**Figure 5:**
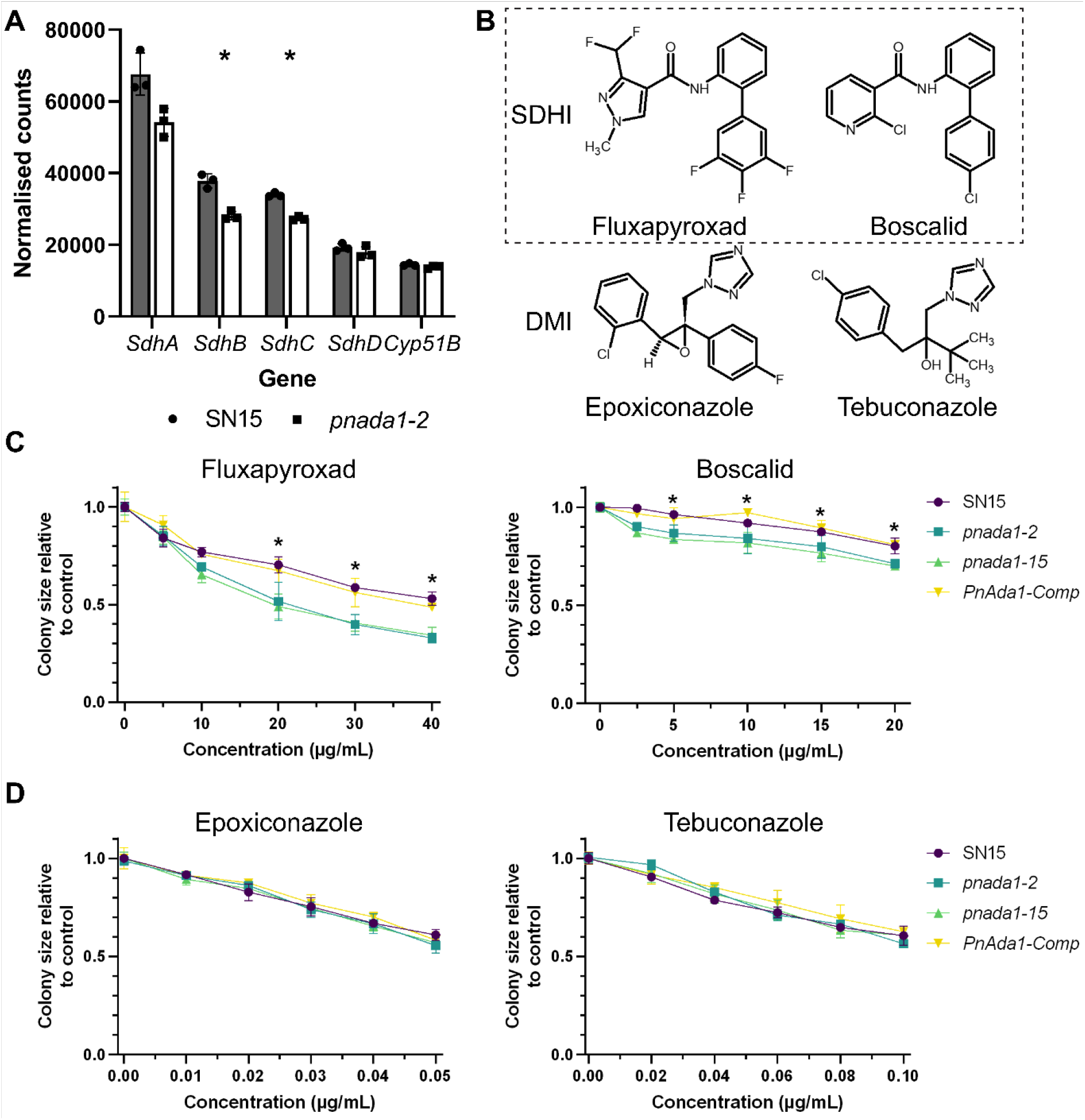
Fungicide sensitivity of *PnAda1* mutants. (**A**) Expression profiles of SDH complex subunit genes and *Cyp51B* between SN15 and *pnada1-2 in vitro*, showing a statistically significant reduction in expression of *SdhB* and *SdhC* but not *SdhA*, *SdhD* or *Cyp51B*. * *p* < 0.01. (**B**) Chemical structures of the tested succinate dehydrogenase inhibitors (SDHI) and demethylase inhibitors (DMI). (**C** and **D**) Growth of fungal strains on V8PDA supplemented with (**C**) the SDHIs fluxapyroxad and boscalid or (**D**) the DMIs epoxiconazole and tebuconazole. Asterisks (*) represent a condition whereby the average colony sizes of SN15 and *PnAda1-Comp* significantly differ from *pnada1-2* and *pnada1-15* according to a two-way ANOVA with Tukey’s HSD (*p* < 0.05).

## DISCUSSION

In this study, a transcriptome-guided approach was used to further characterise the regulatory function of PnAda1 within the PnPf2 network in *P. nodorum*. Integrating *in vitro* and *in planta* comparative RNA sequencing revealed that the deletion of *PnAda1* perturbed the fungal transcriptome leading to a delay in the progression of host infection. This effect is due to the disruption of the initial gene expression needed for host colonisation rather than by directly repressing NE expression. Although the *pnada1-2* mutant expresses NEs at SN15 (or higher) levels, it accumulates less biomass and develops delayed lesions. Based on evidence gathered, we hypothesised that this is due to down-regulation of a large suite of CAZymes required to degrade plant-host tissue and acquire nutrients much like PnPf2 mutants (Jones et al., 2019). When relative infection stage rather than identical/comparable time is considered, the apparent overexpression of NE disappears, indicating that *pnada1-2* is developmentally delayed. Overall, PnAda1 regulates a large set of virulence-associated genes and appears to coordinate the transition from early infection to established necrotrophy, integrating nutrient acquisition, stress responses, and infection-related gene expression to ensure successful disease development.

In SN15, the transcriptome was extensively shifted between the early (3 dpi) and later, more established (7 dpi) stage of infection, whereas this transition was markedly reduced in *pnada1-2*, which at 7 dpi retained a transcriptional profile resembling a SN15 3 dpi-like earlier stage of infection. This included a failure for *pnada1-2* to down-regulate NE genes, secreted cell wall-degrading enzymes and proteases, and genes associated with translation and oxidative phosphorylation. We therefore propose that PnAda1 functions less as a regulator of any individual pathway than as a coordinator of the physiological transition linked to fungal development and infection. This interpretation consistent with both the pleiotropic phenotypes of Ada1 orthologs across fungi (Chen et al., 2020, Gai et al., 2022, Son et al., 2011, Zhao et al., 2022) and the delayed development, rather than significantly reduced, virulence of *PnAda1*-deletion mutants as we had reported previously (John et al., 2024).

The developmentally-reduced virulence of *PnAda1*-deletion mutants appears to be independent of both host penetration and NE-mediated pathogenesis, mirroring the NE-independent virulence defects of *PnPf2* and *PnVeA* deletion mutants (John et al., 2024, Morikawa et al., 2024), but contrasting with the regulator *PnCsn6* (Morikawa et al., 2026), where partial rescue by pre-wounding suggests that PnCsn6 and PnAda1 may govern distinct pathogenicity mechanisms. Nevertheless, our *in planta* data revealed a distinct role for PnAda1 in NE regulation: whereas NE transcripts declined between 3 and 7 dpi in SN15, all tested NE genes remained elevated in *pnada1-2* at 7 dpi. PnAda1 is therefore required not for the activation of NE genes (akin to the role of PnPf2 (Rybak et al., 2017)) but for their timely down-regulation as infection progresses. Such attenuation may help to match effector deployment at the correct stage of infection progression, and its failure in *pnada1-2* is consistent with the delayed disease progression in the absence of *PnAda1*. A comparable timing defect was evident for some secreted degradative machinery, for example; the putative cutinase (*JI435_143050*) was suppressed in *pnada1-2* during early infection yet over-expressed at 7 dpi, reinforcing the view that PnAda1 governs the timing of the infection-associated/activated secretome. As cutinases and secreted proteases contribute to cuticle penetration, apoplastic nutrient acquisition and the degradation of host defence proteins during early infection (Kubicek et al., 2014, Jashni et al., 2015), their reduced expression is consistent with the delayed ability of the PnAda1-deletion mutants to colonise the host compared to SN15.

In parallel to PnAda1 activity described here, the bZIP factor BIP1 of *Magnaporthe oryzae*, which is essential for the establishment of infection but, like PnAda1, is dispensable for host penetration – *BIP1* deletion is not rescued by wounding. Furthermore, BIP1 activates a specific set of early invasion genes encoding effectors, secreted enzymes and secondary-metabolism proteins (Lambou et al., 2024). Although BIP1 acts as an activator and PnAda1 as a more general regulator, both illustrate how a single bZIP factor can gate the deployment and timing of the infection-associated secretome in phytopathogenic fungi.

Beyond development and virulence, the conserved functions of Ada1 remain poorly defined. Functional enrichment implicated PnAda1 in amino acid and nitrogen metabolism, consistent with observations for AaAda1 in *Alternaria alternata* (Gai et al., 2022), and defined growth assays here confirmed a requirement for PnAda1 in the utilisation of several nitrogen and carbon sources. Notably, *pnada1-2* grew poorly on nitrate despite RNA-Seq indicating the up-regulation of the nitrate assimilation gene *NIA1* both *in vitro* and *in planta*, suggesting that downstream nitrate metabolism is perturbed. As urea and nitrate are ultimately assimilated into ammonium (Tudzynski, 2014), the concurrent defect in ammonium utilisation may underlie a broader nitrogen-regulon phenotype. Additionally, the enhanced growth of each *PnAda1-*mutant on media containing sodium acetate and ethanol is consistent with PnAda1 acting to restrain flux through the acetate-assimilation route, since the alcohol dehydrogenase, aldehyde dehydrogenase and acetyl-CoA synthetase-associated genes are all significantly upregulated in our RNA-Seq data for *pnada1-2 in vitro,* and rank among the most abundant transcripts.

The *in planta* RNA-Seq analysis for the *PnAda1-*mutant extend this nitrogen-carbon phenotype into a broader model of nutrient homeostasis. During early infection, *pnada1-2* strongly induced high-affinity nutrient-scavenging systems -- methionine and sulfate acquisition, and ammonium and nitrate assimilation -- consistent with the *PnAda1-*mutant attempting to compensate for the reduced deployment of the extracellular degradative enzymes that normally liberate host-derived nutrients (Divon and Fluhr, 2007). By established infection at 7 dpi, however, these assimilation genes were no longer induced, and the wider transporter repertoire was repressed, coinciding with the impaired *in planta* growth of the *PnAda1*-mutant. Early scavenging giving way to a general contraction of nutrient-acquisition gene expression, suggests that a primary consequence of *PnAda1* loss during infection is a failure to sustain nutrient supply, a determinant for successful host colonisation (Snoeijers et al., 2000, Solomon et al., 2003). As nutrient limitation is also a signal that shapes fungal development and the deployment of virulence factors (Fernandez and Wilson, 2014), such a nutrient-acquisition defect could contribute to the impaired developmental progression and overall reduced virulence in the *PnAda1-*mutant. Framing PnAda1 as a regulator required to maintain nutrient-acquisition capacity therefore offers a parsimonious link between its metabolic, developmental and virulence phenotypes.

*In vitro* transcriptome-guided approach also uncovered a marginal link between PnAda1 and fungicide sensitivity. SDHI resistance typically arises from mutations in the succinate dehydrogenase (Sdh) subunits (Peng et al., 2021, Liu et al., 2023, Lalève et al., 2014), whereas over-expression of a target gene can confer resistance to other classes such as the DMIs (Price et al., 2015). Here we showed *PnAda1*-deletion mutants were hypersensitive to SDHIs but not DMIs. As PnAda1 appears to influence SDHI tolerance specifically rather than fungicide tolerance in general, it may be plausible that mechanistically, it is through the reduced expression of *SdhB* and *SdhC* directly or indirectly. This identifies a transcriptionally encoded determinant of intrinsic SDHI sensitivity in *P. nodorum*. The hypersensitivity of *pnada1-2* to SDHI fungicides is of particular interest, given reduced SdhB and SdhC expression lowers the abundance of the fungicide target complex without directly abolishing respiration, indicating that PnAda1 may represent a candidate for chemical or biological potentiation in combination with current fungicide-based approaches. Such approaches co-targeting or supplementing fungicide treatments with other chemistries have been utilised for plant pathogens, including *P. nodorum* with promising results on overlapping *PnAda1*-related mechanisms. For example, the *PnAda1-* regulated redox-active cell-wall targeting compound thymol has been used successfully as a co-applicant to overcome *P. nodorum* fungicide resistance (Shcherbakova et al., 2021). More direct applications, such as utilising a RNAi-based approach to antagonise *PnAda1* expression, may also prove effective -- as recently demonstrated with the silencing of another efflux regulator TF in the phytopathogen *Botrytis cinerea* (López-Laguna et al., 2026).

The processes influenced by PnAda1; nitrogen metabolism, calcium homeostasis, succinate dehydrogenase function, cell wall and membrane biogenesis, oxidative stress tolerance and virulence, are all closely tied to mitochondrial activity (John et al., 2024, Fung et al., 2025, Li et al., 2014, Li et al., 2020, Carvalho et al., 2020, Qu et al., 2012, Dagley et al., 2011, Shingu-Vazquez and Traven, 2011). The *in planta* de-repression of oxidative phosphorylation and ribosomal genes in *pnada1-2* provides direct transcriptional evidence. KEGG enrichment of heterokaryon-incompatibility genes for *pnada1-2 in vitro* further echoes the cell-cycle role reported for FpAda1 in *Fusarium pseudograminearum* (Chen et al., 2020). One mechanism by which a single regulator could couple these processes is mitochondrial retrograde signalling, whereby the transcriptional response to mitochondrial status adjusts nitrogen and carbon metabolism and stress tolerance (Liu and Butow, 2006). Consistent with this, mitochondrial dysfunction can activate Crz1, driving the expression of transporter and chitin synthase genes and enhancing cell wall biogenesis (Li et al., 2020) -- a phenotype also seen in *PnAda1*-deletion mutants, which additionally de-repress the calcium-signalling regulator PnCrz1 (Choupannejad et al., 2025). As several Ada1 orthologs are also required for oxidative stress tolerance (Chen et al., 2020, Gai et al., 2022, Zhao et al., 2022, John et al., 2024), we speculate that a conserved link between Ada1 and mitochondrial function may underlie the pleiotropy of this regulator. In future, direct measurements of respiration and mitochondrial membrane potential would provide a valuable test of this hypothesis.

Taken together, this study positions PnAda1 as a broad coordinator of the transcriptional programmes underlying *P. nodorum* development and infection, with roles spanning nitrogen metabolism, abiotic-stress and fungicide tolerance, and the temporal control of the infection transcriptome, including NE genes. By reframing the pleiotropy of Ada1 as the coordination of developmental transitions -- potentially through a conserved association with mitochondrial function -- this work provides new insights for dissecting the roles of Ada1 orthologs across fungal pathogens.

## MATERIALS AND METHODS

### Strain and cultures

*P. nodorum* strains used in this study are outlined in **Supplemental Table S3** and maintained on V8 potato dextrose agar (V8PDA) (150 mL L^−1^ Campbell’s V8 Juice, 3 g L^−1^ CaCO_3_, 10 g L^−1^ Difco PDA and 10 g L^−1^ agar) as described previously (Solomon et al., 2004).

### RNA extraction, sequencing and quality control

RNA sequences were prepared as previously described (Morikawa et al., 2024). For *in vitro* samples, SN15 and *pnada1-2* were grown on V8PDA for 12 days and bulk RNA was extracted using TRIzol™ reagent (Invitrogen) and subsequently treated with DNase I (New England Biolabs), following manufacturers’ protocols. *In planta* RNA was harvested from infected wheat leaves as described previously (Verdonk et al., 2025). RNA-Seq library preparation and sequencing were performed by the Australian Genome Research Facility (https://www.agrf.org.au/) (*in vitro*) or by GENEWIZ Azenta (https://www.genewiz.com) (*in planta*), with 150 bp paired-end stranded sequences generated on Illumina NovaSeq platforms 6000 S4 (*in vitro*) or NovaSeq X (*in planta*). The experiment was performed in biological triplicate (*in vitro*) or quadruplicate (*in planta*). The quality of the raw RNA-Seq reads was assessed in FastQC v0.11.9 (https://www.bioinformatics.babraham.ac.uk/projects/fastqc/), with sequencing adapters trimmed using fastp v1.0.1 (Chen et al., 2018) and filtered to retain sequencing reads with a quality score of Q30 or above.

### Comparative RNA-Sequencing and statistical analyses

The trimmed paired reads were mapped to the *P. nodorum* SN15 genome (Bertazzoni et al., 2021) using HISAT2 (Kim et al., 2019) with default settings. The outputs were sorted and converted using SAMtools (Li et al., 2009). The sorted mapped reads were counted using featureCounts v2.0.0 from the Subread package v2.0.2 (Liao et al., 2014).

Only genes with more than five mapped reads were retained for further analysis. The following analyses were performed on R v4.1.2 (https://www.R-project.org/): differentially expressed genes (DEGs) were determined using DESeq2 v1.34.0 with Log_2_ fold change (LFC) shrinkage applied to reduce noise from genes with low counts (Love et al., 2014, Zhu et al., 2019). The criteria for DEGs were defined as the LFC against the null hypothesis −0.58 ≤ LFC ≤ 0.58 (i.e. *H_a_*: |LFC| ≥ 0.58), with a Benjamini-Hochberg correction adjusted *p*-value significance threshold of 0.01. The cutoffs |LFC| ≥ 1 and |LFC| ≥ 2 were also analysed to compare the effects of various LFC cutoff values. Similarly, the adjusted *p-*value significance thresholds of 0.05 and 0.001 were also analysed. Differences between medians of average normalised gene expression were performed using the Mann-Whitney U test (two comparisons) or the Kruskal-Wallis test with *post-hoc* Dunn’s test (more than two comparisons) with Benjamini-Hochberg correction unless otherwise specified.

### Determination of functional enrichment in differentially expressed genes

Up- and down-regulated genes were analysed separately for functional enrichment, as it increases sensitivity in detecting unique enriched functions (Hong et al., 2014). However, the analysis was performed with total DEGs if the separation did not yield unique enriched functions. Gene Ontology (GO) terms were assigned to *P. nodorum* genes using InterProScan v5.76 (Jones et al., 2014). The R package GOseq v1.26.0 (Young et al., 2010) was used to find overrepresented GO terms in DEGs. ShinyGO v0.85 (Ge et al., 2020) was utilised for the Kyoto Encyclopedia of Genes and Genomes (KEGG) pathway enrichment analysis with the species set to “*Parastagonospora nodorum* STRINGdb”.

### Phylogenetic and sequence analyses

Sequence orthologs of succinate dehydrogenase enzymes in *P. nodorum* SN15 were identified using TBLASTN (Camacho et al., 2009) with the species restricted to *P. nodorum* and using the representative SdhA, SdhB, SdhC and SdhD sequences from a previous study as queries (Mair et al., 2016).

Analysis of gene promoters for enriched DNA motifs were conducted as described previously (Jones et al., 2019, John et al., 2024). Briefly, a 1500-bp upstream region (or to the next annotated gene) of predicted start codons of DE genes were used in MEME Suite (Bailey et al., 2015) v5.5.9 to identify putative motifs. Position weight matrices for top non-redundant *de novo* motifs were analysed with FIMO (Grant et al., 2011) and searched against known motifs with Tomtom (Gupta et al., 2007). STREME (Bailey, 2021) was used for enriched PnPf2 M1 (Jones et al., 2019) and M2 motifs (John et al., 2024), as well as common bZIP TF core ACGT-like (Foster et al., 1994) motifs (ACGT, TGACGTCA, TGACGT, CGTCA, RTGACGTCAY) or TRE/AP-1-like (Fernandes et al., 1997, Paluh et al., 1988) motifs (TGACTCA, TGASTCA, TGACTAA, TTACTAA, TTAGTAA).

### Nitrogen and carbon metabolism, stress tolerance and fungicide sensitivity assays

For the nitrogen and carbon metabolism assays, *P. nodorum* strains were grown for 12 days on MM agar (Solomon et al., 2004) with varying nitrogen sources (24 mM glutamine, ammonium chloride, lysine, leucine, sodium nitrate, proline or urea) or carbon sources (24 mM sucrose, starch, mannitol, sodium acetate and ethanol). The colony sizes were normalised to the colonies grown on MM with the primary nitrogen source glutamine or primary carbon source sucrose to account for the reduced growth of *PnAda1*-deletion mutants.

For the stress tolerance assays, *P. nodorum* strains were grown for seven days on MM agar with 24 mM glutamine as a nitrogen source supplemented with either 10-20 mM calcium chloride, 10-100 µg/mL sodium dodecyl sulphate, 5 µg/mL Congo Red or 5-15 µg/mL Calcofluor White (Sigma-Aldrich)(Morikawa et al., 2026). The colony sizes were normalised to the control colonies grown on MM agar without the addition of an abiotic stressor.

For the fungicide sensitivity assays, 1 × 1 mm agar plugs of *P. nodorum* strains were inoculated on 2 mL V8PDA in a 12-well cell culture plate in biological triplicates supplemented with either 5-40 µg/mL fluxapyroxad, 2.5-20 µg/mL boscalid, 0.01-0.05 µg/mL epoxiconazole or 0.02-0.1 µg/mL tebuconazole. Colony sizes were measured five days post-inoculation. The colony sizes were normalised to the SN15 control colonies grown without the addition of a fungicide. The chemical structures of the tested fungicides were drawn using ChemSketch Freeware (https://www.acdlabs.com/resources/free-chemistry-software-apps/chemsketch-freeware/).

### Wheat infection assays

Detached leaf assay (DLA) was performed as described previously (Solomon et al., 2004, Verdonk et al., 2025). The *P. nodorum* mycelial/hyphal mixture was homogenised using a TissueLyser II (QIAGEN) with a 1-mm diameter steel bead at 30 Hz for 10 sec to prepare 50 mg/mL of hyphae in 0.02% (v/v) Tween 20 solution. The mycelial/hyphal mixture was spread using a paintbrush that had been pre-wet with 0.02% (v/v) Tween 20 solution onto approximately 4 cm of the first wheat leaves (cv. Halberd), which were embedded adaxially into benzimidazole agar (75 mg/L benzimidazole and 15 g/L agar). For (pre-)wounded leaves, wheat leaves were pierced through with a hypodermic 25G needle (Interpath) immediately prior to mycelial inoculation. The infection was allowed to develop for 3-, 5-, 7- and 10-days post inoculation. Negative treatments in all assays were performed with 0.02% (v/v) Tween 20 solution only. NE gene expression assays were performed via qPCR as previously described (Rybak et al., 2017, Morikawa et al., 2024) using primers listed in **Supplemental Table S4**.

## DISCLOSURE OF AI USE

The authors used AI to improve the clarity and readability of the manuscript and subsequently reviewed and edited all content.

## ACKNOWLEDGEMENTS

This study was conducted by the Centre for Crop and Disease Management, a co-investment between the Grains Research and Development Corporation (GRDC) and Curtin University – grant CUR1403-002BLX. The funders had no role in the experimental design, data collection and analysis, the decision to publish, or the preparation of the manuscript. The authors declare that there are no conflicts of interest.

The authors thank Dr. Evan John and David Jiang for sample preparation and preliminary analysis.

## CRediT

<u>Shota Morikawa:</u> Conceptualisation, Data Curation, Formal analysis, Investigation, Methodology, Writing - original draft. <u>Leon Lenzo:</u> Data Curation, Formal analysis, Investigation. <u>Keshara Colomba Thanthrige:</u> Investigation. <u>Steven Chang:</u> Methodology, Validation. <u>Kar-Chun Tan:</u> Conceptualisation, Funding acquisition, Project administration, Resources, Supervision, Writing - review & editing. <u>Callum Verdonk:</u> Conceptualisation, Data Curation, Formal analysis, Investigation, Methodology, Supervision, Validation, Writing - original draft, Writing - review & editing.

## DATA AVAILABILITY

The raw RNA sequences used in this study are available in the National Center for Biotechnology Information Sequence Read Archive database under the BioProject ID PRJNA1509083.

## SUPPLEMENTARY MATERIAL CAPTIONS

**Supplemental Figure S1**: Analysis of comparative *in vitro* RNA-Seq between SN15 and *pnada1-2*. (**A**) Scatter plots with average normalised counts of genes in SN15 on the x-axis and in *pnada1-2* on the y-axis. The right scatter plot shows the plot zoomed in at 200000 normalised counts for both axes. The displayed region is marked with a blue dotted box in the left scatter plot. Log_2_ fold change (LFC) cutoffs are displayed at the top and right of the scatterplots as straight lines, indicating the divergence of high LFC cutoffs from the adjusted p-value at higher basal gene expression levels. Areas are colour-coded according to the LFC cutoffs in (**B**) and (**C**). (**B, C** and **D**) Boxplots of average normalised counts in SN15 with different LFC cutoffs in *pnada1-2* showing that high LFC cutoffs disproportionately filter highly expressed genes. Comparisons made with up-regulated (**B**) and down-regulated (**C**) genes in *pnada1-2* (adj. *p* < 0.01). (**D**) Boxplots of average normalised counts in SN15 with different adjusted *p*-value cutoffs in *pnada1-2* showing that high stringent adjusted *p*-value cutoffs disproportionately filter lowly expressed genes. “n” is the number of genes and “x̃” represents the median normalised count. The y-axis is capped at 20000 normalised counts. Asterisks represent statistically significant difference between the medians using the Kruskal-Wallis test followed by the Dunn’s test with Benjamini-Hochberg correction (adj. *p* < 0.05(*); < 0.0001(****)). “n.s.” signifies “not significant” (i.e. adj. *p* > 0.05).

**Supplemental Figure S2**: Average normalised counts of genes in enriched Gene Ontology (GO) terms in SN15 and *pnada1-2 in vitro*. The x-axis is capped at 20000 normalised counts. Asterisks represent statistically. “n.s.” signifies no significant difference between the medians using the Kruskal-Wallis test followed by the Dunn’s test with Benjamini-Hochberg correction.

**Supplemental Figure S3:** Relative *in planta Parastagonospora nodorum* pathogen RNA-Seq transcript counts (*n* = 4 for each contrast). Left panel shows the percentage (%) of *P. nodorum* reads during wheat host infection at 3 dpi and 7 dpi for *pnada1-2* and SN15. Right panel shows the total read count (in millions) of *P. nodorum* at each timepoint (contrast) for both *pnada1-2* and SN15.

**Supplemental Figure S4:** Representative vegetative morphologies of *P. nodorum* strains grown on defined minimal media (MM) agar with (**A**) various nitrogen sources (NH_4_Cl, ammonium chloride; NaNO_3_, sodium nitrate) or (**B**) various carbon sources (NaOAc, sodium acetate). Measured colony diameter values are shown in **Figure 4**.

**Supplemental Table S1 (separate .xlsx file):** RNA-Seq analysis differentially expressed genes in *pnada1-2* relative to SN15 for *in vitro*, *in planta* 3 dpi and *in planta* 7dpi. Sheets:

*In vitro:* DE_pnada1-2_iv
*In planta* 3 dpi: DE_pnada1-2_ip_3dpi
*In planta* 7 dpi: DE_pnada1-2_ip_7dpi

**Supplemental Table S2 (separate .xlsx file):** RNA-Seq analysis differentially expressed genes in *pnada1-2* for all characterised genes from *P. nodorum* SN15. Statistically significant genes **bolded**. LFC and adjusted *p-*values (p-adj) shown for each gene.

**Supplemental Table S3**: *P. nodorum* strains used in this study.

**Supplemental Table S4**: Primers used in this study.

## REFERENCES

Amoutzias, G. D., Veron, A. S., Weiner, J., 3rd, Robinson-Rechavi, M., Bornberg-Bauer, E., Oliver, S. G. & Robertson, D. L. 2007. One billion years of bzip transcription factor evolution: conservation and change in dimerization and DNA-binding site specificity. Molecular Biology and Evolution, 24, 827–35.

Bailey, T. L. 2021. Streme: accurate and versatile sequence motif discovery. Bioinformatics, 37, 2834–2840.

Bailey, T. L., Johnson, J., Grant, C. E. & Noble, W. S. 2015. The MEME Suite. Nucleic Acids Research, 43, W39–W49.

Bertazzoni, S., Jones, D. A. B., Phan, H. T., Tan, K.-C. & Hane, J. K. 2021. Chromosome-level genome assembly and manually-curated proteome of model necrotroph *Parastagonospora nodorum* SN15 reveals a genome-wide trove of candidate effector homologs, and redundancy of virulence-related functions within an accessory chromosome. BMC Genomics, 22, 382.

Camacho, C., Coulouris, G., Avagyan, V., Ma, N., Papadopoulos, J., Bealer, K. & Madden, T. L. 2009. Blast+: architecture and applications. BMC Bioinformatics, 10, 421.

Carvalho, E. J., Stathopulos, P. B. & Madesh, M. 2020. Regulation of Ca(2+) exchanges and signaling in mitochondria. Current Opinion in Physiology, 17, 197–206.

Chen, L., Ma, Y., Zhao, J., Geng, X., Chen, W., Ding, S., Li, H. & Li, H. 2020. The bZIP transcription factor FpAda1 is essential for fungal growth and conidiation in *Fusarium pseudograminearum*. Current Genetics, 66, 507–515.

Chen, S., Zhou, Y., Chen, Y. & Gu, J. 2018. fastp: an ultra-fast all-in-one FASTQ preprocessor. Bioinformatics, 34, i884–i890.

Choupannejad, R., Sharifnabi, B., Collemare, J., Gholami, J. & Mehrabi, R. 2025. The candidate transcription factors PnAtfa, PnCrz1, and PnVf19 contribute to fungal morphogenesis, abiotic stress tolerance, and pathogenicity in the wheat pathogen *Parastagonospora nodorum*. Fungal Biology, 129, 101565.

Cutler, S. B. & Caten, C. E. 1999. Characterisation of the nitrite reductase gene (NII1) and the nitrate-assimilation gene cluster of *Stagonospora* (*Septoria*) *nodorum*. Current Genetics, 36, 282–9.

Dagley, M. J., Gentle, I. E., Beilharz, T. H., Pettolino, F. A., Djordjevic, J. T., Lo, T. L., Uwamahoro, N., Rupasinghe, T., Tull, D. L., Mcconville, M., Beaurepaire, C., Nantel, A., Lithgow, T., Mitchell, A. P. & Traven, A. 2011. Cell wall integrity is linked to mitochondria and phospholipid homeostasis in *Candida albicans* through the activity of the post-transcriptional regulator Ccr4-Pop2. Molecular Microbiology, 79, 968–89.

Divon, H. H. & Fluhr, R. 2007. Nutrition acquisition strategies during fungal infection of plants. FEMS Microbiology Letters, 266, 65–74.

Duarte Hospital, C., Tête, A., Debizet, K., Imler, J., Tomkiewicz-Raulet, C., Blanc, E. B., Barouki, R., Coumoul, X. & Bortoli, S. 2023. SDHi fungicides: An example of mitotoxic pesticides targeting the succinate dehydrogenase complex. Environment International, 180, 108219.

Fernandes, L., Rodrigues-Pousada, C. & Struhl, K. 1997. Yap, a novel family of eight bzip proteins in *Saccharomyces cerevisiae* with distinct biological functions. Molecular and Cellular Biology, 17, 6982–93.

Fernandez, J. & Wilson, R. A. 2014. Cells in cells: morphogenetic and metabolic strategies conditioning rice infection by the blast fungus *Magnaporthe oryzae*. Protoplasma, 251, 37–47.

Foster, R., Izawa, T. & Chua, N.-H. 1994. Plant bZIP proteins gather at ACGT elements. The FASEB Journal, 8, 192–200.

Fung, T. S., Ryu, K. W. & Thompson, C. B. 2025. Arginine: at the crossroads of nitrogen metabolism. The EMBO Journal, 44, 1275–1293.

Gai, Y., Li, L., Liu, B., Ma, H., Chen, Y., Zheng, F., Sun, X., Wang, M., Jiao, C. & Li, H. 2022. Distinct and essential roles of bZIP transcription factors in the stress response and pathogenesis in *Alternaria alternata*. Microbiology Research, 256, 126915.

Ge, S. X., Jung, D. & Yao, R. 2020. Shinygo: a graphical gene-set enrichment tool for animals and plants. Bioinformatics, 36, 2628–2629.

Geng, Q., Li, H., Wang, D., Sheng, R. C., Zhu, H., Klosterman, S. J., Subbarao, K. V., Chen, J. Y., Chen, F. M. & Zhang, D. D. 2022. The *Verticillium dahliae* Spt-Ada-Gcn5 Acetyltransferase Complex Subunit Ada1 Is Essential for Conidia and Microsclerotia Production and Contributes to Virulence. Frontiers in Microbiology, 13, 852571.

Grant, C. E., Bailey, T. L. & Noble, W. S. 2011. Fimo: scanning for occurrences of a given motif. Bioinformatics, 27, 1017–1018.

Gupta, S., Stamatoyannopoulos, J. A., Bailey, T. L. & Noble, W. S. 2007. Quantifying similarity between motifs. Genome Biology, 8, R24.

Hong, G., Zhang, W., Li, H., Shen, X. & Guo, Z. 2014. Separate enrichment analysis of pathways for up- and downregulated genes. Journal of the Royal Society Interface, 11, 20130950.

Howard, K., Foster, S. G., Cooley, R. N. & Caten, C. E. 1999. Disruption, replacement, and cosuppression of nitrate assimilation genes in *Stagonospora nodorum*. Fungal Genetics and Biology, 26, 152–62.

Ipcho, S. V., Hane, J. K., Antoni, E. A., Ahren, D., Henrissat, B., Friesen, T. L., Solomon, P. S. & Oliver, R. P. 2012. Transcriptome analysis of *Stagonospora nodorum*: gene models, effectors, metabolism and pantothenate dispensability. Molecular Plant Pathology, 13, 531–45.

Jashni, M. K., Mehrabi, R., Collemare, J., Mesarich, C. H. & De Wit, P. J. 2015. The battle in the apoplast: further insights into the roles of proteases and their inhibitors in plant-pathogen interactions. Frontiers in Plant Science, 6, 584.

John, E., Singh, K. B., Oliver, R. P. & Tan, K. C. 2021. Transcription factor control of virulence in phytopathogenic fungi. Molecular Plant Pathology, 22, 858–881.

John, E., Verdonk, C., Singh, K. B., Oliver, R. P., Lenzo, L., Morikawa, S., Soyer, J. L., Muria-Gonzalez, J., Soo, D., Mousley, C., Jacques, S. & Tan, K.-C. 2024. Regulatory insight for a Zn2Cys6 transcription factor controlling effector-mediated virulence in a fungal pathogen of wheat. PLOS Pathogens, 20, e1012536.

Jones, D. A. B., John, E., Rybak, K., Phan, H. T. T., Singh, K. B., Lin, S. Y., Solomon, P. S., Oliver, R. P. & Tan, K. C. 2019. A specific fungal transcription factor controls effector gene expression and orchestrates the establishment of the necrotrophic pathogen lifestyle on wheat. Scientific Reports, 9, 15884.

Jones, P., Binns, D., Chang, H.-Y., Fraser, M., Li, W., Mcanulla, C., Mcwilliam, H., Maslen, J., Mitchell, A., Nuka, G., Pesseat, S., Quinn, A. F., Sangrador-Vegas, A., Scheremetjew, M., Yong, S.-Y., Lopez, R. & Hunter, S. 2014. InterProScan 5: genome-scale protein function classification. Bioinformatics, 30, 1236–1240.

Kim, D., Paggi, J. M., Park, C., Bennett, C. & Salzberg, S. L. 2019. Graph-based genome alignment and genotyping with HISAT2 and HISAT-genotype. Nature Biotechnology, 37, 907–915.

Kong, S., Park, S.-Y. & Lee, Y.-H. 2015. Systematic characterization of the bZIP transcription factor gene family in the rice blast fungus, *Magnaporthe oryzae*. Environmental Microbiology, 17, 1425–1443.

Kubicek, C. P., Starr, T. L. & Glass, N. L. 2014. Plant cell wall-degrading enzymes and their secretion in plant-pathogenic fungi. Annual Review of Phytopathology, 52, 427–51.

Lalève, A., Gamet, S., Walker, A. S., Debieu, D., Toquin, V. & Fillinger, S. 2014. Site-directed mutagenesis of the P225, N230 and H272 residues of succinate dehydrogenase subunit B from *Botrytis cinerea* highlights different roles in enzyme activity and inhibitor binding. Environmental Microbiology, 16, 2253–66.

Lambou, K., Tag, A., Lassagne, A., Collemare, J., Clergeot, P. H., Barbisan, C., Perret, P., Tharreau, D., Millazo, J., Chartier, E., De Vries, R. P., Hirsch, J., Morel, J. B., Beffa, R., Kroj, T., Thomas, T. & Lebrun, M. H. 2024. The bZIP transcription factor BIP1 of the rice blast fungus is essential for infection and regulates a specific set of appressorium genes. PLOS Pathogens, 20, e1011945.

Li, H., Handsaker, B., Wysoker, A., Fennell, T., Ruan, J., Homer, N., Marth, G., Abecasis, G. & Durbin, R. 2009. The Sequence Alignment/Map format and SAMtools. Bioinformatics, 25, 2078–9.

Li, L., Hu, X., Xia, Y., Xiao, G., Zheng, P. & Wang, C. 2014. Linkage of oxidative stress and mitochondrial dysfunctions to spontaneous culture degeneration in *Aspergillus nidulans*. Molecular and Cellular Proteomics, 13, 449–61.

Li, Y., Zhang, Y., Zhang, C., Wang, H., Wei, X., Chen, P. & Lu, L. 2020. Mitochondrial dysfunctions trigger the calcium signaling-dependent fungal multidrug resistance. Proceedings of the National Academy of Sciences, 117, 1711–1721.

Liao, Y., Smyth, G. K. & Shi, W. 2014. featureCounts: an efficient general purpose program for assigning sequence reads to genomic features. Bioinformatics, 30, 923–30.

Liu, Y., Sun, Y., Bai, Y., Cheng, X., Li, H., Chen, X. & Chen, Y. 2023. Study on Mechanisms of Resistance to SDHI Fungicide Pydiflumetofen in *Fusarium fujikuroi*. Journal of Agricultural and Food Chemistry, 71, 14330–14341.

Liu, Z. & Butow, R. A. 2006. Mitochondrial retrograde signaling. Annual Review of Genetics, 40, 159–85.

López-Laguna, A., Mota-Maldonado, V., Morales, Y., Perez-Garcia, A. & Fernandez-Ortuno, D. 2026. Spray-induced gene silencing targeting the transcription factor Bcmrr1: A sustainable RNAi-based strategy to control *Botrytis cinerea* and overcome multidrug resistance. Plant Disease.

Love, M. I., Huber, W. & Anders, S. 2014. Moderated estimation of fold change and dispersion for RNA-seq data with DESeq2. Genome Biology, 15, 550.

Mair, W., Lopez-Ruiz, F., Stammler, G., Clark, W., Burnett, F., Hollomon, D., Ishii, H., Thind, T. S., Brown, J. K., Fraaije, B., Cools, H., Shaw, M., Fillinger, S., Walker, A. S., Mellado, E., Schnabel, G., Mehl, A. & Oliver, R. P. 2016. Proposal for a unified nomenclature for target-site mutations associated with resistance to fungicides. Pest Management Science, 72, 1449–59.

Mcdonald, M. C., Williams, S. J. & Solomon, P. S. 2023. The Role of Tox Effector Proteins in the *Parastagonospora Nodorum*–Wheat Interaction. In: Scott, B. & Mesarich, C. (eds.) Plant Relationships: Fungal-Plant Interactions. Cham: Springer International Publishing.

Morikawa, S., Herbst, C., John, E., Croll, D., Mousley, C., Henares, B., Tan, K.-C. & Verdonk, C. 2026. A Conserved Transcription Factor Domain Drives Necrotrophic Effector-Mediated Virulence and Putative Protein Interactions in *Parastagonospora nodorum*. Molecular Plant-Microbe Interactions.

Morikawa, S., Verdonk, C., John, E., Lenzo, L., Sbaraini, N., Turo, C., Li, H., Jiang, D., Chooi, Y.-H. & Tan, K.-C. 2024. The Velvet transcription factor PnVea regulates necrotrophic effectors and secondary metabolism in the wheat pathogen *Parastagonospora nodorum*. BMC Microbiology, 24, 299.

Paluh, J. L., Orbach, M. J., Legerton, T. L. & Yanofsky, C. 1988. The cross-pathway control gene of *Neurospora crassa*, cpc-1, encodes a protein similar to GCN4 of yeast and the DNA-binding domain of the oncogene v-jun-encoded protein. Proceedings of the National Academy of Sciences, 85, 3728–3732.

Peng, J., Sang, H., Proffer, T. J., Gleason, J., Outwater, C. A., Jung, G. & Sundin, G. W. 2021. A Method for the Examination of SDHI Fungicide Resistance Mechanisms in Phytopathogenic Fungi Using a Heterologous Expression System in *Sclerotinia sclerotiorum*. Phytopathology, 111, 819–830.

Price, C. L., Parker, J. E., Warrilow, A. G., Kelly, D. E. & Kelly, S. L. 2015. Azole fungicides - understanding resistance mechanisms in agricultural fungal pathogens. Pest Management Science, 71, 1054–8.

Qu, Y., Jelicic, B., Pettolino, F., Perry, A., Lo, T. L., Hewitt, V. L., Bantun, F., Beilharz, T. H., Peleg, A. Y., Lithgow, T., Djordjevic, J. T. & Traven, A. 2012. Mitochondrial sorting and assembly machinery subunit Sam37 in *Candida albicans*: insight into the roles of mitochondria in fitness, cell wall integrity, and virulence. Eukaryotic Cell, 11, 532–44.

Richards, J. K., Kariyawasam, G. K., Seneviratne, S., Wyatt, N. A., Xu, S. S., Liu, Z., Faris, J. D. & Friesen, T. L. 2022. A triple threat: the *Parastagonospora nodorum* SnTox267 effector exploits three distinct host genetic factors to cause disease in wheat. New Phytologist, 233, 427–442.

Rybak, K., See, P. T., Phan, H. T., Syme, R. A., Moffat, C. S., Oliver, R. P. & Tan, K. C. 2017. A functionally conserved Zn2Cys6 binuclear cluster transcription factor class regulates necrotrophic effector gene expression and host-specific virulence of two major Pleosporales fungal pathogens of wheat. Molecular Plant Pathology, 18, 420–434.

Seshasayee, A. S., Sivaraman, K. & Luscombe, N. M. 2011. An overview of prokaryotic transcription factors : a summary of function and occurrence in bacterial genomes. Subcellular Biochemistry, 52, 7–23.

Shcherbakova, L., Mikityuk, O., Arslanova, L., Stakheev, A., Erokhin, D., Zavriev, S. & Dzhavakhiya, V. 2021. Studying the Ability of Thymol to Improve Fungicidal Effects of Tebuconazole and Difenoconazole Against Some Plant Pathogenic Fungi in Seed or Foliar Treatments. Frontiers in Microbiology, Volume 12 - 2021.

Shingu-Vazquez, M. & Traven, A. 2011. Mitochondria and fungal pathogenesis: drug tolerance, virulence, and potential for antifungal therapy. Eukaryotic Cell, 10, 1376–83.

Snoeijers, S. S., Pérez-García, A., Joosten, M. H. A. J. & De Wit, P. J. G. M. 2000. The Effect of Nitrogen on Disease Development and Gene Expression in Bacterial and Fungal Plant Pathogens. European Journal of Plant Pathology, 106, 493–506.

Solomon, P. S., Lee, R. C., Wilson, T. J. G. & Oliver, R. P. 2004. Pathogenicity of *Stagonospora nodorum* requires malate synthase. Molecular Microbiology, 53, 1065–1073.

Solomon, P. S., Tan, K.-C. & Oliver, R. P. 2003. The nutrient supply of pathogenic fungi; a fertile field for study. Molecular Plant Pathology, 4, 203–210.

Son, H., Seo, Y. S., Min, K., Park, A. R., Lee, J., Jin, J. M., Lin, Y., Cao, P., Hong, S. Y., Kim, E. K., Lee, S. H., Cho, A., Lee, S., Kim, M. G., Kim, Y., Kim, J. E., Kim, J. C., Choi, G. J., Yun, S. H., Lim, J. Y., Kim, M., Lee, Y. H., Choi, Y. D. & Lee, Y. W. 2011. A phenome-based functional analysis of transcription factors in the cereal head blight fungus, *Fusarium graminearum*. Plos Pathogens, 7, e1002310.

Tian, C., Li, J. & Glass, N. L. 2011. Exploring the bzip transcription factor regulatory network in *Neurospora crassa*. Microbiology, 157, 747–759.

Tudzynski, B. 2014. Nitrogen regulation of fungal secondary metabolism in fungi. Frontiers in Microbiology, Volume 5–2014.

Verdonk, C., Screaigh, I., Morikawa, S., Lenzo, L., Mousley, C. & Tan, K.-C. 2025. Dissecting the role of early infection-expressed putative transcription factors in the pathogenicity of *Parastagonospora nodorum* on wheat. Access Microbiology, 7.

Weirauch, M. T. & Hughes, T. R. 2011. A catalogue of eukaryotic transcription factor types, their evolutionary origin, and species distribution. Subcellular Biochemistry, 52, 25–73.

Young, M. D., Wakefield, M. J., Smyth, G. K. & Oshlack, A. 2010. Gene ontology analysis for Rna-seq: accounting for selection bias. Genome Biology, 11, R14.

Zhao, Q., Pei, H., Zhou, X., Zhao, K., Yu, M., Han, G., Fan, J. & Tao, F. 2022. Systematic Characterization of bzip Transcription Factors Required for Development and Aflatoxin Generation by High-Throughput Gene Knockout in *Aspergillus flavus*. Journal of Fungi, 8.

Zhu, A., Ibrahim, J. G. & Love, M. I. 2019. Heavy-tailed prior distributions for sequence count data: removing the noise and preserving large differences. Bioinformatics, 35, 2084–2092.

Zulak, K. G., Chang, S., Tan, K.-C., Turo, C., Oliver, R. P. & Lopez-Ruiz, F. J. 2025. Dissecting allele-specific fungicide resistance mechanisms by heterologous expression of the demethylase inhibitor target gene *Cyp51* in a phytopathogen model. bioRxiv, 2025.10.16.682737.

