## Supplementary File for "The bZIP transcription factor PnAda1 functions as a regulator of virulence, fungicide tolerance and necrotrophy in the wheat pathogen *Parastagonospora nodorum*"

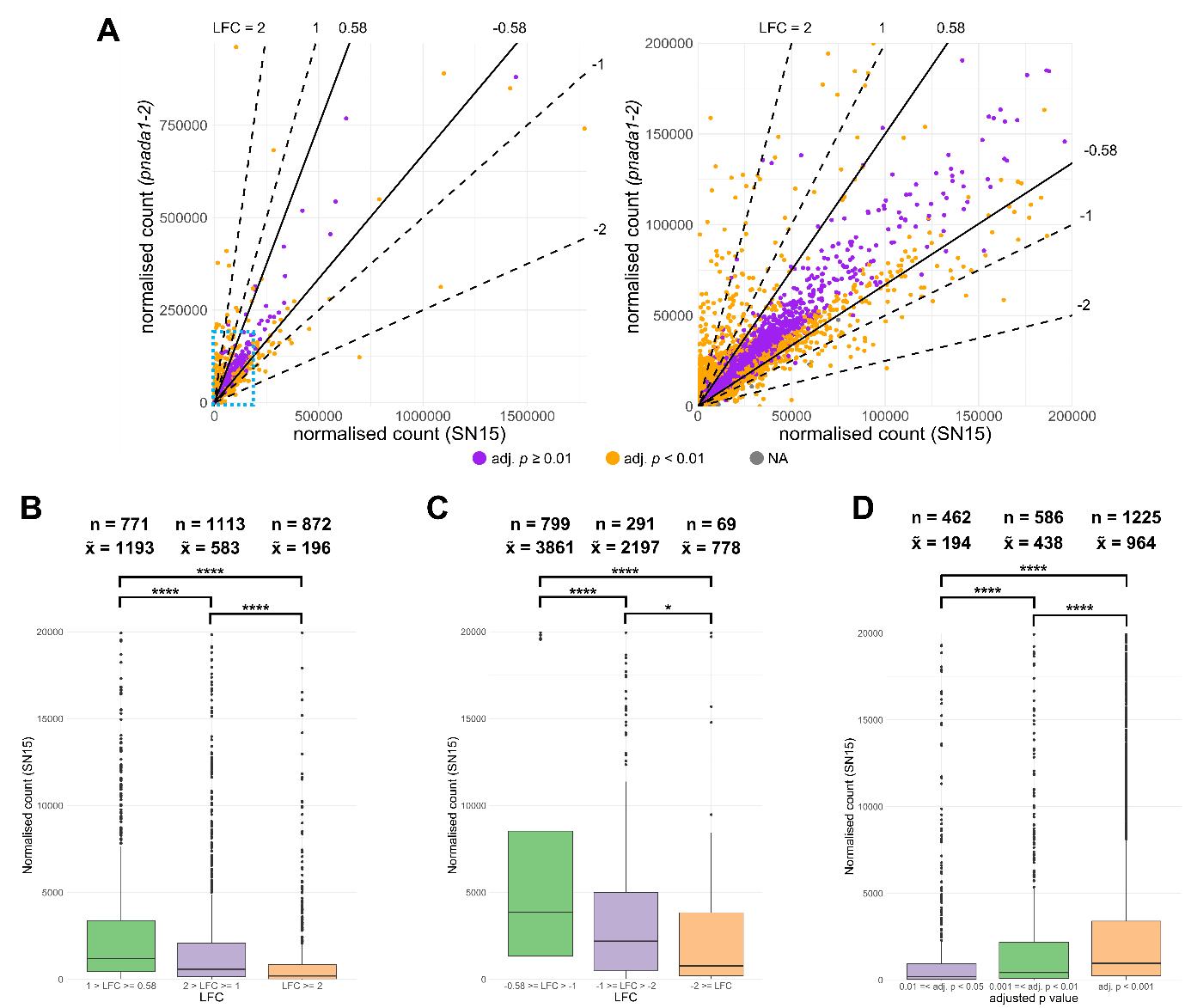

**Supplemental Figure S1**: Analysis of comparative *in vitro* RNA-Seq between SN15 and *pnada1-2*. (**A**) Scatter plots with average normalised counts of genes in SN15 on the x-axis and in *pnada1-2* on the y-axis. The right scatter plot shows the plot zoomed in at 200000 normalised counts for both axes. The displayed region is marked with a blue dotted box in the left scatter plot. Log_2_ fold change (LFC) cutoffs are displayed at the top and right of the scatterplots as straight lines, indicating the divergence of high LFC cutoffs from the adjusted p-value at higher basal gene expression levels. Areas are colour-coded according to the LFC cutoffs in (**B**) and (**C**). (**B, C** and **D**) Boxplots of average normalised counts in SN15 with different LFC cutoffs in *pnada1-2* showing that high LFC cutoffs disproportionately filter highly expressed genes. Comparisons made with up-regulated (**B**) and down-regulated (**C**) genes in *pnada1-2* (adj. *p* < 0.01). (**D**) Boxplots of average normalised counts in SN15 with different adjusted *p*-value cutoffs in *pnada1-2* showing that high stringent adjusted *p*-value cutoffs disproportionately filter lowly expressed genes. “n” is the number of genes and “x**̃**” represents the median normalised count. The y-axis is capped at 20000 normalised counts. Asterisks represent statistically significant difference between the medians using the Kruskal-Wallis test followed by the Dunn’s test with Benjamini-Hochberg correction (adj. *p* < 0.05(*); < 0.0001(****)). “n.s.” signifies “not significant” (i.e. adj. *p* > 0.05).

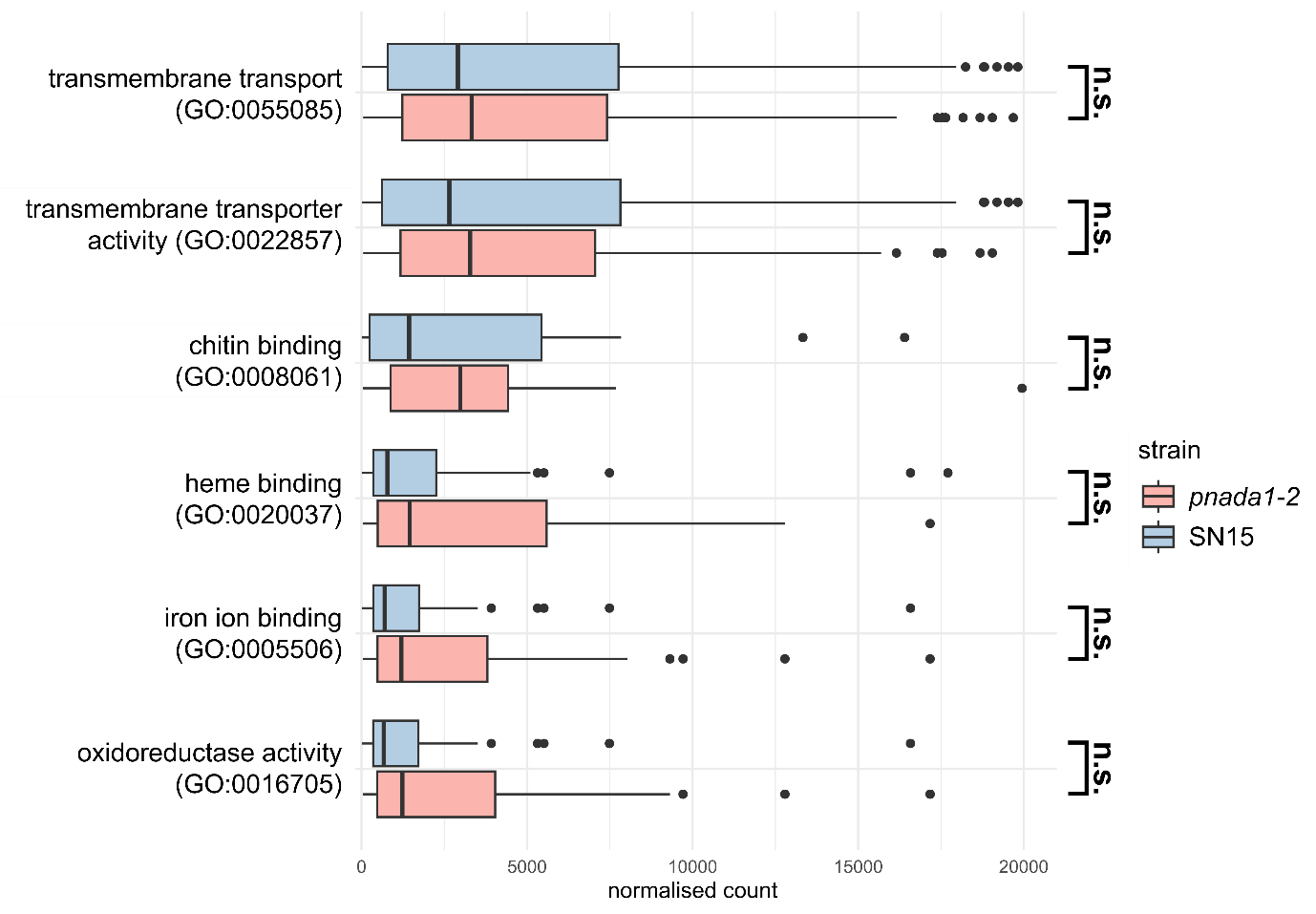

**Supplemental Figure S2**: Average normalised counts of genes in enriched Gene Ontology (GO) terms in SN15 and *pnada1-2 in vitro*. The x-axis is capped at 20000 normalised counts. Asterisks represent statistically. “n.s.” signifies no significant difference between the medians using the Kruskal-Wallis test followed by the Dunn’s test with Benjamini-Hochberg correction.

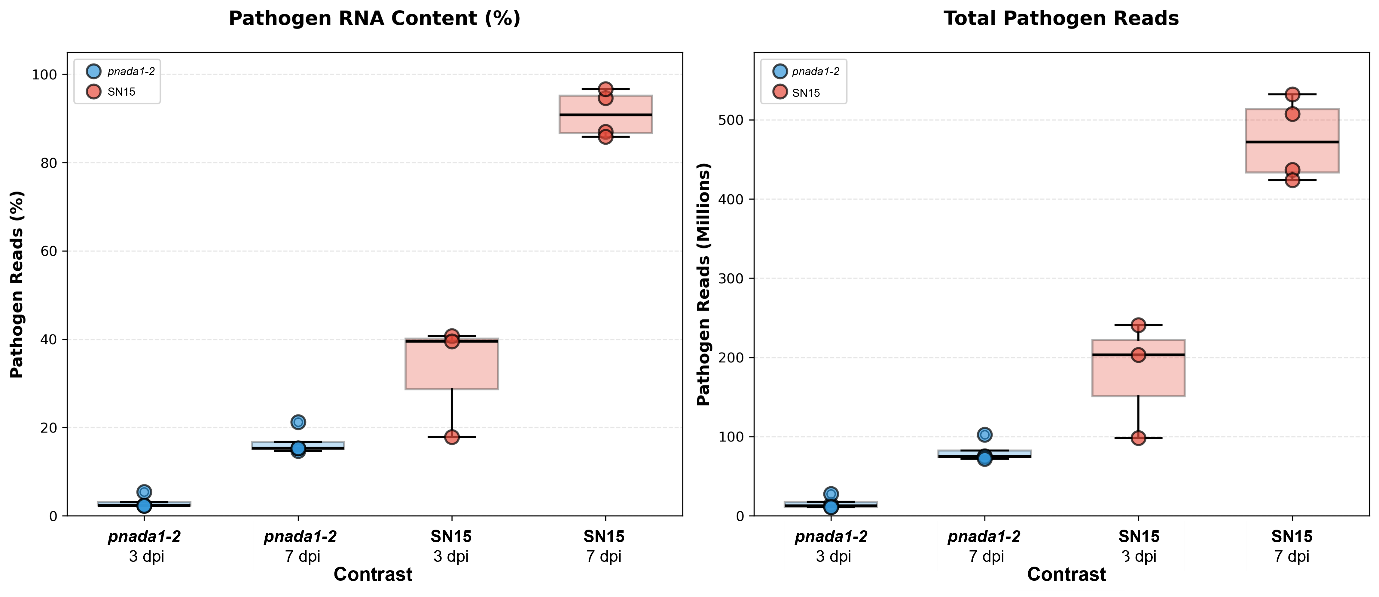

**Supplemental Figure S3:** Relative *in planta* *Parastagonospora nodorum* pathogen RNA-Seq transcript counts (*n* = 4 for each contrast). Left panel shows the percentage (%) of *P. nodorum* reads during wheat host infection at 3 dpi and 7 dpi for *pnada1-2* and SN15. Right panel shows the total read count (in millions) of *P. nodorum* at each timepoint (contrast) for both *pnada1-2* and SN15.

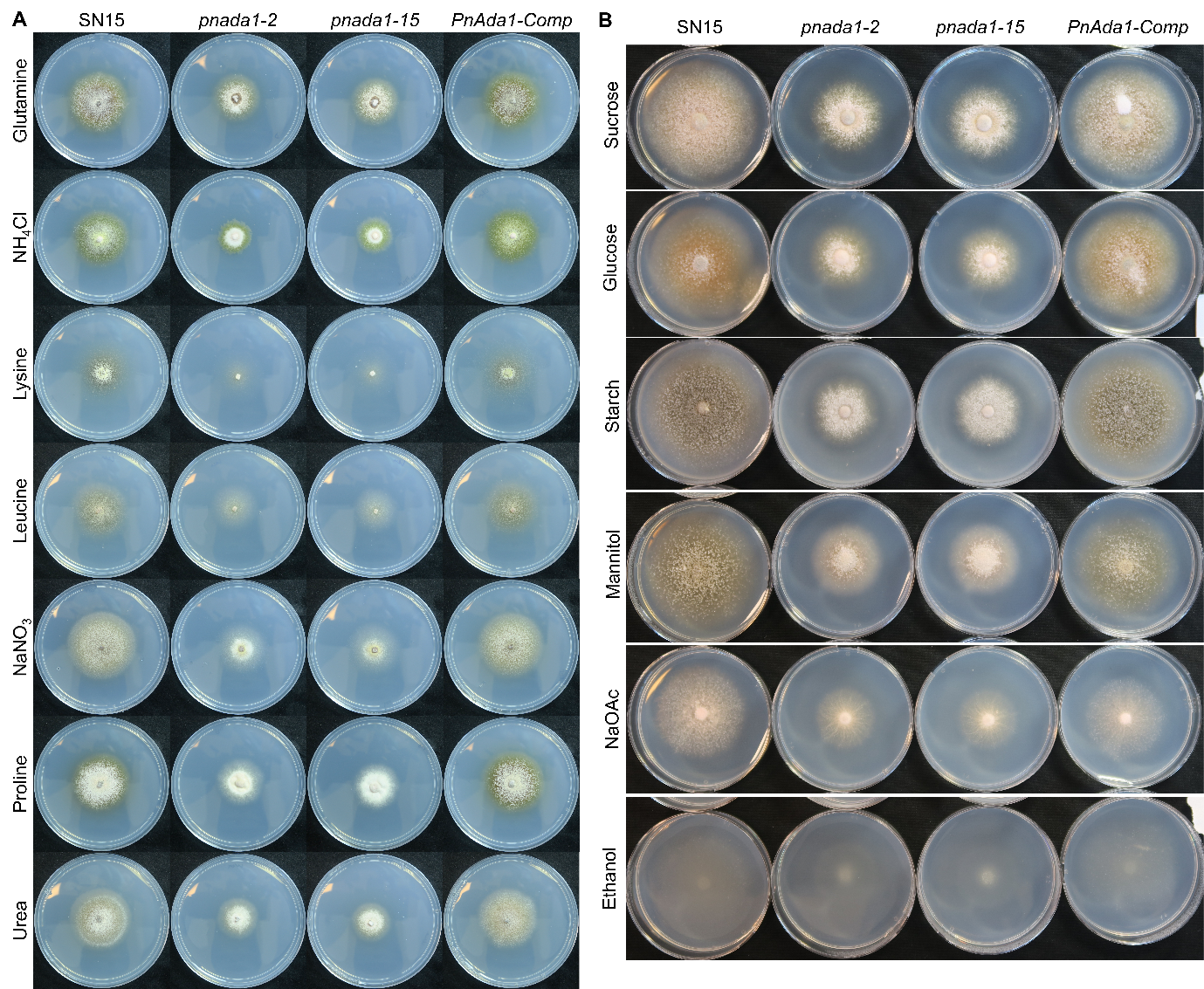

**Supplemental Figure S4:** Representative vegetative morphologies of *P. nodorum* strains grown on defined minimal media (MM) agar with (**A**) various nitrogen sources (NH_4_Cl, ammonium chloride; NaNO_3_, sodium nitrate) or (**B**) various carbon sources (NaOAc, sodium acetate). Measured colony diameter values are shown in **Figure 4**.

**Supplemental Table S1 (separate .xlsx file):** RNA-Seq analysis differentially expressed genes in *pnada1-2* relative to SN15 for *in vitro*, *in planta* 3 dpi and *in planta* 7dpi. Sheets:

*In vitro:* DE_pnada1-2_iv

*In planta* 3 dpi: DE_pnada1-2_ip_3dpi

*In planta* 7 dpi: DE_pnada1-2_ip_7dpi

**Supplemental Table S2 (separate .xlsx file):** RNA-Seq analysis differentially expressed genes in *pnada1-2* for all characterised genes from *P. nodorum* SN15. Statistically significant genes **bolded**. LFC and adjusted *p-*values (p-adj) shown for each gene.

**Supplemental Table S3**: *P. nodorum* strains used in this study.

| **Strain** | **Description** | **Source** |
| --- | --- | --- |
| SN15 | *Parastagonospora nodorum* wildtype | Department of Primary Industries and Regional Development, Western Australia |
| *pnada1-2* | SN15 with an in-frame deletion of *PnAda1* (*JI435_044860*) integrated into *PnAda1* locus, variant 2. Hyg^R^ | (9) |
| *pnada1-15* | SN15 with an in-frame deletion of *PnAda1* (*JI435_044860*) integrated into *PnAda1* locus, variant 15. Hyg^R^ | (9) |
| *PnAda1-Comp* | *pnada1-2* with a complementation of *PnAda1* randomly integrated into the genome. | (9) |

**Supplemental Table S4**: Primers used in this study.

| **Name** | **Sequence** | **Purpose** |
| --- | --- | --- |
| ToxA_qPCR_F | CGATCCCGGTTACGAAATC | qPCR NEs |
| ToxA_qPCR_R | TTGACATGCAGCTTCCCTG | qPCR NEs |
| Tox1_qPCR_F | TGGTCTTGTCAGTAGCCTTTGC | qPCR NEs |
| Tox1_qPCR_R | TCCTGGAGTATGGCAAATTG | qPCR NEs |
| Tox3_qPCR_F | AATGTCGACCGTTTTGACC | qPCR NEs |
| Tox3_qPCR_R | GGTTGCCGCAGTTGATATAA | qPCR NEs |
| Actin_qPCR_F | AGTCGAAGCGTGGTATCCT | qPCR relative marker *Act1* |
| Actin_qPCR_R | ACTTGGGGTTGATGGGAG | qPCR relative marker *Act1* |
